# Bidirectional regulation of postmitotic H3K27me3 distributions underlie cerebellar granule neuron maturation dynamics

**DOI:** 10.1101/2022.10.10.511582

**Authors:** Vijyendra Ramesh, Fang Liu, Melyssa S. Minto, Urann Chan, Anne E. West

## Abstract

The functional maturation of neurons is a prolonged process that extends well beyond mitotic exit and is mediated by the chromatin-dependent orchestration of gene transcription programs. We find that the postnatal maturation of cerebellar granule neurons (CGNs) requires dynamic changes in the genomic distribution of histone H3 lysine 27 trimethylation (H3K27me3), demonstrating a function for this chromatin modification beyond its role in cell fate specification. The developmental loss of H3K27me3 at promoters of genes that turn on as CGNs mature is facilitated by the lysine demethylase, and ASD-risk gene, Kdm6b. Interestingly, inhibition of the H3K27 methyltransferase EZH2 in newborn CGNs not only blocks the repression of progenitor genes but also impairs the induction of mature CGN genes, showing the importance of bidirectional H3K27me3 regulation across the genome. These data demonstrate that H3K27me3 turnover in developing postmitotic neurons regulates the temporal coordination of gene expression programs that underlie functional neuronal maturation.

## Introduction

Cell identity is determined during differentiation by the coordinated actions of chromatin regulatory factors, which serve to establish the accessibility of gene regulatory elements that control transcription of cell-type specific genes. Histone proteins play a key role in determining the architecture of chromatin, and specific post-translational modifications of the histones are tightly associated with the transcriptional state of a given locus or gene [1]. Among these modifications, histone methylation occurs on lysine and arginine residues of specific histones, and has roles in both gene activation and repression [2], and acts to alter gene transcription by recruiting the site-specific binding of effector proteins that carry out specialized functions [3]. The trimethylation of lysine 27 on histone H3 (H3K27me3) is a repressive histone modification that is best known for its role during cell-fate determination [4]. H3K27 methylation is deposited by the Polycomb repressive complex 2 (PRC2) lysine methyltransferases EZH1 and EZH2 [5, 6]. H3K27me3 can be enzymatically removed by the H3K27-specific lysine demethylases KDM6A (UTX) and KDM6B (JMJD3) [7]. In mice, null mutations in *Ezh2* result in early embryonic lethality [5], *Kdm6a* knockouts show severely impaired mid-gestational development [8], and *Kdm6b* knockouts die at birth [9] suggesting the physiological importance of H3K27me3 regulation for mammalian embryonic development.

Neurons are born very early in their overall lifespan, and they undergo significant transcriptional and functional changes during postmitotic stages of their developmental maturation. The enzymes that write and erase H3K27me3 remain expressed in neurons both during postnatal stages of development and in the adult brain [10, 11], suggesting that they have functions beyond fate determination that remain to be understood. Conditional deletion of both *Ezh1* and *Ezh2* in neurons of the adult mouse causes a slow global loss of H3K27me3 as well as progressive neurodegeneration and death showing the importance of these enzymes in adult neurons [12]. One possibility is that H3K27me3 is required for fate maintenance. For example, deleting EED in developing postmitotic dopamine or serotonin neurons *in vivo* leads not only to loss of H3K27me3 but also to reduced expression of tyrosine hydroxylase (TH) and tryptophan hydroxylase 2 (TPH2), respectively, which mediate dopamine and serotonin synthesis to define the transmitter phenotype of these neuromodulatory neurons [13]. Other studies suggest that H3K27me3 might be required in terminally differentiated neurons to persistently repress genes that have the potential to drive trans-differentiation to alternate cell fates [12, 14].

The global depletion of H3K27me3 in postmitotic neurons takes months to appear after deletion of EZH1/2 [12] revealing that this histone modification is remarkably stable across the bulk of the genome. Although histone replacement is a major mechanism of H3K27me3 removal in dividing cells [15], histone turnover is very slow in postmitotic neurons [16]. Interestingly, H3K27-selective lysine demethylases are widely expressed in the adult brain suggesting that these enzymes may have a particularly important role in the local regulation of histone methylation patterns in postmitotic cells [17]. Both *Kdm6a* and *Kdm6b* are expressed in neurons over the course of brain development, though only *Kdm6b* shows enhanced expression following exposure of cells to neuronal differentiation cues [11, 18–20]. Germline knockout of *Kdm6b* does not impair the gross architecture of the embryonic brain, but it does result in perinatal lethality caused by disruption of the functional maturation of a respiratory circuit that is required for proper breathing after birth [9]. This phenotype arises as a result of the lost demethylase function of Kdm6b, because survival cannot be rescued by transgenic expression of a demethylase-dead Kdm6b in the knockout mice [9]. These data are important because they suggest that demethylation mediated by Kdm6b is essential for the proper function of postmitotic neurons. Notably, *KDM6B* has been identified as a high confidence autism spectrum disorder (ASD) risk gene in humans [21], and *Kdm6b* haploinsufficiency in mice is associated with ASD-like impairments in sociability [22]. However the underlying chromatin and transcriptional mechanisms of these Kdm6b mutation phenotypes remain to be elucidated.

To begin to identify functions of histone H3K27 demethylation in postmitotic stages of neurodevelopment, we previously conditionally deleted *Kdm6a* and *Kdm6b* in *Atoh1*-expressing cerebellar granule neuron (CGN) progenitors of the mouse. We used CGNs to model neurodevelopment because they comprise more than 99% of all neurons [23] and 85% of cells [24] in the cerebellum, enabling the *in vivo* comparison of chromatin across CGN developmental states with limited contamination from other cell types. When purified and placed in *ex vivo* culture, CGN progenitors coordinately exit the cell cycle, minimizing heterogeneity of cell developmental stage across the population [25]. In both preparations, gene transcription programs change over time coincident with changes in CGN biology (*e.g*., proliferation, migration, synapse formation, synapse maturation [26, 27]). Loss of *Kdm6a/b* had no effect on postnatal morphogenesis of the cerebellum, demonstrating these enzymes are not required *in vivo* for regulation of CGN progenitor proliferation. However knockdown of *Kdm6b* in cultured CGNs impaired the developmental induction of genes during the period of postmitotic neuronal maturation [11]. Based on these data, we hypothesized that H3K27me3 may serve as a temporal gatekeeper of these maturation genes, delaying their activation beyond the immediate period of terminal differentiation to the neuronal fate.

Here to directly test the hypothesis that H3K27me3 regulation serves as a mechanism of neuronal maturation we determined the role of H3K27me3 dynamics in changing programs of gene transcription in postmitotic CGNs. We used ChIP-seq and a CUT&RUN time course study to identify the genomic location and timing of differential sites of H3K27me3 enrichment across the genome of fate-committed CGNs developing *in vivo* and in culture. We observed that H3K27me3 distributions continue to change even after fate committed CGN progenitors exit the cell cycle, and that the loss of H3K27me3 at gene promoters is associated with the activation of genes induced as neurons mature. We used both genetic and pharmacological approaches to disrupt the action of H3K27me3 regulatory enzymes and show evidence of roles for both the lysine demethylase KDM6B and the lysine methyltransferase EZH2 in the induction of a postmitotic program of gene transcription that underlies functional maturation of CGNs. Taken together, our data advance understanding of how programs of gene expression are temporally orchestrated by changing H3K27me3 distributions in the chromatin of maturing neurons.

## Materials and Methods

### Animal Husbandry

We performed all procedures under an approved protocol from the Duke University Institutional Animal Care and Use Committee. *Kdm6b* floxed mice described in [28] were obtained from Jackson Labs (Stock #029615, RRID #IMSR_JAX:029615). These mice contain *loxP* sites flanking exon 14-20, encoding the catalytic Jumonji-C domain, of the mouse *Kdm6b* gene on chromosome 11. *Atoh1-cre* mice were obtained from Jackson Labs (Stock #011104, RRID #IMSR_JAX:011104). Mice were genotyped by ear clipping at the time of weaning using protocols described on the Jackson Labs website. CD-1 IGS mice (#Strain 022, #IMSR_CRL:022) and C57BL/6NCrl mice (#Strain 027, RRID #IMSR_CRL:027) were obtained from Charles River Laboratories. Both male and female mice were used for all experiments in this study.

### CGN Culture

Cerebella of CD1-IGS mice were dissected at P6-P8. Cells were dissociated and run on a Percoll gradient to isolate cerebellar granule neuron progenitors (GNPs) as previously described [29]. GNPs were plated at a density of 1 million cells/well in a 24-well plate for RNA or protein isolation. At day *in vitro* 2 (DIV2) CGNs were treated with 1 μM Cytosine Arabinoside (AraC) (Sigma #C1768) to block division of any non-neuronal cells. KDM6 and EZH2 inhibitors were added at the doses and days described in the text.

### RT-qPCR

CGNs were lysed in TRIzol Reagent (Thermo #15596026) at a volume of 0.25 mL per 1 million cells for RNA purification. Isolated RNA was analyzed using a nanodrop to determine concentration and purity, after which it was treated with DNase I (NEB #M0303S). We made Oligo(dT) primed cDNA using SuperscriptII Reverse Transcriptase (Invitrogen #19064014). qPCR was carried out on an Applied Biosystems Quantstudio 3 Thermal cycler using Power SYBR Green PCR Master Mix (Thermo #4367659) and PCR primers targeting the gene/locus of interest (**Supplementary Information, Table S1**).

### RNA-seq

RNA for RNA-seq was obtained by processing cerebellar tissue or CGN cultures with Trizol reagent followed by clean up on the Zymo Direct-zol RNA miniprep kit (Zymo #R2052). RNA purity was measured to ensure samples had A260/280 and A260/230 values > 1.9. RNA was then polyA enriched and 150 bp paired-end sequencing was performed by Novogene, Inc on an Illumina Hi-seq 2000 machine.

### Chromatin Immunoprecipitation

#### Cerebellar Tissue

We pooled cerebellum from three C57BL/6NCrl (Charles River Laboratories IMSR Cat# CRL:027, RRID: IMSR_CRL:027) P7 mice, two P14 mice and one P60 mouse for each biological replicate. Both male and female mice were used. Cerebellum samples were dounced in a glass homogenizer containing 2 ml of 1% formaldehyde (w/v) PBS buffer per sample and kept at 25 °C for 15 min, washed twice with cold PBS, lysed in 600 μl lysis buffer (1% SDS (w/v), 10 mM EDTA, and 50 mM Tris, pH 8.1). The cross-linked material was sonicated with a Bioruptor (Diagenode), with output set to “high” with 30 s on/off cycles to an average size range of 150–350 bp which was then determined by agarose gel electrophoresis. Sonicated chromatin was centrifuged for 10 minutes at 14000 RPM at 4°C, after which the supernatant was diluted 10-fold in dilution buffer (0.01% SDS, 1.1% Triton X-100, 1.2 mM EDTA, 16.7 mM Tris-HCl, pH 8.1, 167 mM NaCl). Prior to immunoprecipitation, 6 μl of antibody (anti-H3K27Me3, Active Motif 39155, RRID: AB_2561020) was incubated with 100 μl of Dynabeads Protein G (Invitrogen 10004D) in PBS for 4-6h at 4 °C. The antibody-bead conjugate was then washed twice with cold PBS, and added to 6 ml of cell lysis for overnight immunoprecipitation at 4°C. The following day, bead-bound DNA-protein complexes were washed, eluted, and purified using a PCR purification kit (Cat# 28104, Qiagen). ChIP-seq libraries were made using the MicroPlex Library Preparation kit V2 (C05010012, Diagenode); and 50 bp single-end sequencing was performed at the Duke Sequencing and Analysis Core Resource on a Hi-Seq 2000 machine. Three independent biological replicates were performed for each antibody and developmental time point and ChIP reagents were prepared according to the Millipore 17–295 ChIP kit.

### Nuclear Isolation of CGNs for CUT&RUN

CGNs were scraped into 1X DPBS at 0.25 mL per 1 million cells and then ‘pop-spun’ according to the REAP method [30], until the rotor reached 7000 rpm. Pellets were then washed once with 1X DPBS and pop-spun again, after which they were resuspended in Nuclei Isolation Buffer (20 mM HEPES pH 7.9, 10 mM KCl, 2 mM Spermidine, 0.1% v/v Triton X-100, 20% v/v glycerol), incubated on ice for 5 minutes and then spun at 2,000g for 5 min at 4°C. After this step the supernatant was removed, and pelleted nuclei were then resuspended in Nuclei Storage Buffer (20 mM Tris-HCl pH 8.0, 75 mM NaCl, 0.5 mM EDTA, 50% v/v glycerol, 1 mM DTT, 0.1 mM PMSF) and stored in - 80°C until ready to process.

### CUT&RUN

CUT&RUN was performed on nuclei isolated from CGN cultures using the CUTANA ChIC/CUT&RUN kit (EpiCypher #14-1408) as per manufacturer guidelines. Specific changes made to the protocol are noted here. Nuclei in Nuclei Storage Buffer were pelleted and resuspended in Nuclei Isolation Buffer. Nuclei were then incubated with activated ConA beads. Antibodies used for CUT&RUN were: H3K27me3 (Active Motif 39155, RRID: AB_2561020), H3K4me3 and IgG (positive and negative controls included in kit). CUT&RUN libraries were made using the NEB Ultra II DNA Library Prep Kit for Illumina (NEB #E7645L), and NEBNext Multiplex Oligos for Illumina (96 Unique Dual Index Primer Pairs) (NEB #E6440S). Library cleanup was performed prior to and after PCR amplification using 0.8X Kapa Hyperpure beads (Roche #08963851001). PCR amplification was performed with the following parameters as described in the EpiCypher CUT&RUN kit: 1) 98°C, 45 sec; 2) 98°C, 15 sec; 60°C, 10 sec x 14 cycles; 3) 72°C, 60 sec. Libraries were then pooled and 50 bp paired-end sequencing was performed at the Duke Sequencing and Analysis Core Resource on a NovaSeq 6000 S-Prime flow cell.

### Sequencing Data Analysis

#### ChlP-seq

ChIP-seq reads were quality scored and screened through fastQC, trimmed for adapter sequences using Trimmomatic 0.38 and then aligned to the mouse GRCm38.p4 reference genome using STAR 2.7.2b. Alignments were filtered for mapped reads using samtools and then normalized using the bamCoverage function of the deepTools2.0 suite. ChIP-seq tracks were generated using bamCoverage, to generate continuous BigWig files. Peak calling was performed using Macs2, with parameters for broad peaks and an FDR threshold of 0.01. The R package DESeq2 [31] was used for read-count normalization and differential binding analysis. R package ChIPseeker [32] was used to annotate ChIP-seq peaks using a window of TSS ±3000. ‘Promoters’ is a group of ChIPseeker terms ‘Promoter (<=1kb), Promoter (1-2 kb) and Promoter (2-3 kb). Gene Body is a group of terms 5’ UTR, 3’ UTR, Intron and Exon.

#### CUT&RUN

CUT&RUN reads were processed the same as ChIP-seq reads until peak-calling. Peak calling was performed using Macs2, with parameters for broad peaks and an FDR threshold of 0.1. Metagene plots were created using SeqPlots [33], by inputting bigwig and BED files and setting a window of TSS +/- 5000 bp.

### RNA-seq

RNA-seq reads were processed similarly to ChIP-seq reads until and excluding peak calling. Read counts for genes were generated using HTseq2 count, with feature type set to ‘exon’, and the Gencode GRCm38.p4 GTF annotation file. DESeq2 was used for read-count normalization and differential expression analysis.

### Data Processing

Heatmaps were generated using the Broad Institute’s Morpheus tool: https://software.broadinstitute.org/Morpheus, with the following clustering parameters: hierarchical clustering, complete linkage, one minus spearman rank correlation, cluster by rows. VST-transformed counts used to generate heatmaps and clusters are provided in Table S2. Gene ontology analysis was performed using the GO consortium GO tool [34, 35] using terms related to ‘biological process’.

### Western Blot

#### Cerebellar Tissue

For preparation of protein lysates from cerebellar tissue, we pooled cerebellum from two P7 mice, one P14 and one P60 mouse for each biological replicate. Cerebellum samples were dounced 50 times with PBS in glass homogenizer, centrifuged at 8000 x g for 5 minutes, and washed twice by cold PBS, then pellets were re-suspended in 300 μL extraction buffer (10 mM HEPES, pH 7.9, 1.5 mM MgCl_2_, 10 mM KCl), supplemented with 1X protease inhibitor cocktail (Roche, Indianapolis, IN). Thereafter, 60μL of 1N HCl was added to the cell lysates and kept on ice for 30 minutes, following which the cell lysates were centrifuged at 11,000 x g for 10 minutes at 4 ºC. Protein concentration of acid-soluble supernatants was measured using Pierce BCA protein assay kit (Cat # 23227, Thermo). 20 μg of acid-soluble supernatant was used for western blots. Samples were loaded onto wells of a 10-20% gradient gel and run at 150V for 30-40 min in running buffer (25 mM Tris-HCl, 192 mM Glycine, 0.1% w/v SDS) after which protein was transferred onto a PVDF membrane (BioRad #1620177) in transfer buffer (25 mM Tris-HCl, 192 mM Glycine, 20% v/v Methanol) for 1 hour at 100V or overnight at 30V. The membrane was blocked for 1 hour in 5% (w/v) BSA (Sigma #A3059) in TBST. After the blocking step, the membrane was incubated with primary antibody overnight at 4°C. The following day, primary antibody was removed, after which the membranes were washed three times with TBST for 5 minutes each, incubated with secondary antibody for 1 hour, and then again washed three times with TBST for 5 minutes each. At the end of this, membranes were imaged using the Li-Cor Odyssey using appropriate exposure times for the target antibodies.

#### CGN Culture

Cells were scraped into PBS from 6 well plates at the relevant endpoints, centrifuged at 200 x g for 5 minutes, and washed twice with PBS. Pellets were then resuspended in 100 μL of extraction buffer (described above), after which 20 μL of 1N HCl was added. Samples were hereafter processed as described above.

#### Antibodies

The antibodies used were H3K27Ac (Abcam ab4729), H3K27Me3 (Active Motif 39155, RRID: AB_2561020), and Total H3 (Tissue: Millipore 05-499, Culture: Cell Signaling 9715). Western blot signals were detected by Li-Cor Odyssey InfraRed Imaging System (Lincoln, NE) with the secondary antibodies, CF770 goat anti-Rabbit, (Biotium 20078) and CF 680 goat anti-mouse, (Biotium 20065). Protein bands were quantified using ImageJ.

### GEO Accession Codes

H3K27me3 ChIP-seq data for P7, P14 and P60 cerebellum, WT and *Kdm6b-cKO* cerebellum; RNA-seq data for WT and *Kdm6b-cKO* cerebellum; CUT&RUN and RNA-seq data for cultured CGNs can be accessed at GEO: GSE212441. RNA-seq data for P7, P14 and P60 cerebellum and DIV0 and DIV7 CGNs, H3K27ac, ZIC1/2 ChIP-seq data and DHS-seq data for P7 and P60 cerebellum were adapted from Frank et. al 2015 [27].

## Author Contributions

Conceptualization: V.R., F.L. and A.E.W.; Methodology: V.R., F.L., U.C., and A.E.W.; Formal Analysis: V.R., M.M.; Investigation V.R.; Resources, A.E.W.; Writing: V.R., A.E.W.; Funding Acquisition: A.E.W.

## Acknowledgements

Fig schematics in Fig 1A, 3A and 5A were created using biorender.com. This work was supported by NIH grant R01NS0988804.

## Results

### Cerebellar Maturation Involves Dynamic, Genome-Wide Changes in H3K27me3 Distribution

To measure H3K27me3 dynamics during cerebellar maturation *in vivo*, we harvested cerebellar cortex from C57BL/6NCrl mice at postnatal days 7, 14, and 60 (Fig. 1A). At P7, about half of all cells in the mouse cerebellum are proliferating granule neuron precursors (GNPs) whereas the rest are immature, postmitotic cerebellar granule neurons (CGNs). By P14 GNP cell division has ceased, and the cerebellum is composed of immature postmitotic CGNs that are migrating to the inner granule layer (IGL) where they receive synapses and fully mature by P60 [24, 27]. Western blot of acid extracted histones showed a significant global increase in total H3K27me3 levels from P7 to P14 that was maintained at P60 (Fig 1B), whereas total H3K27ac fell in a similar temporal pattern (Fig. S1A). These data are consistent with the PRC2-dependent restriction of cell potential upon terminal differentiation [4].

**Fig 1.**
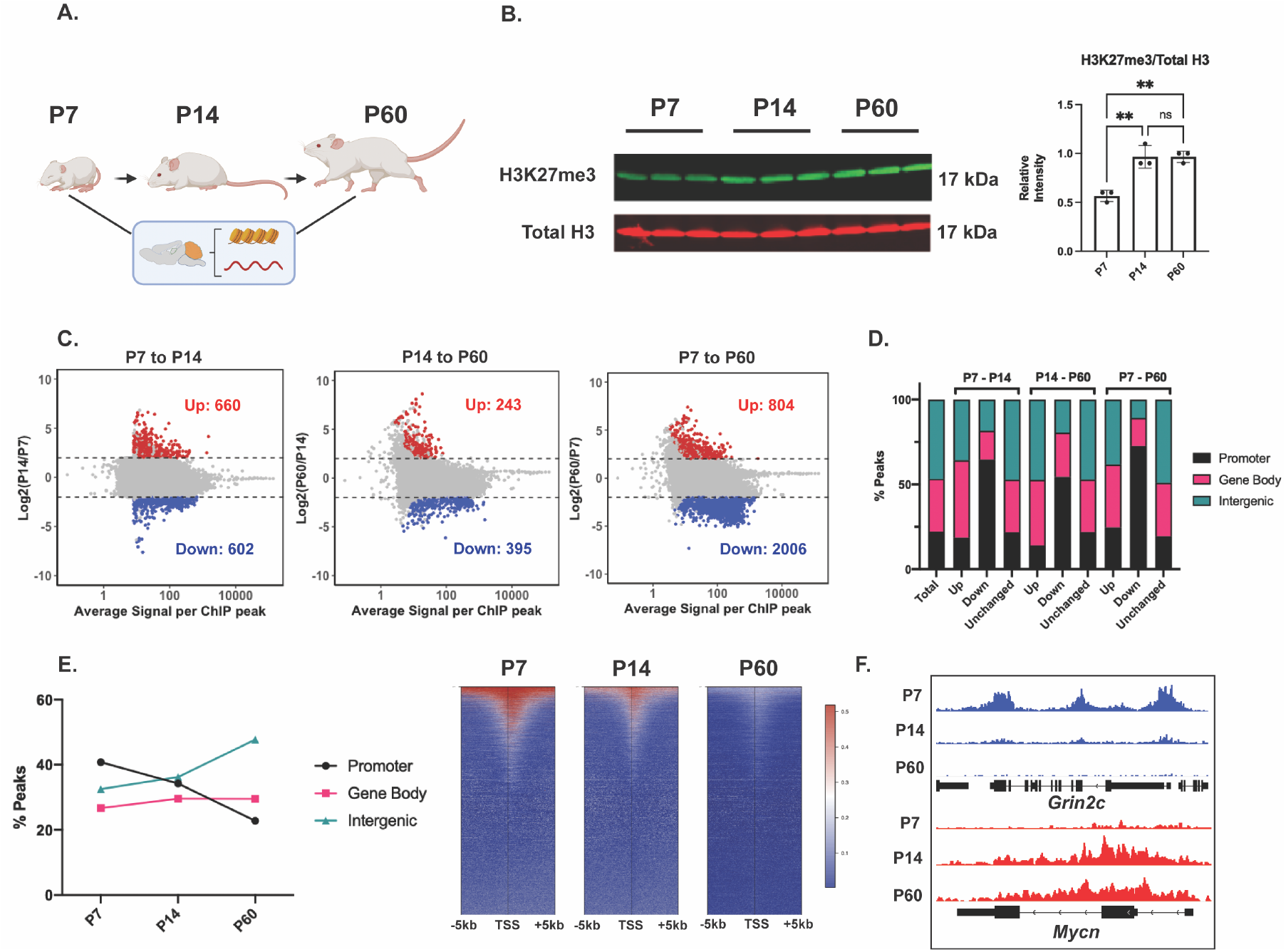
The maturing mouse cerebellum undergoes a genome-wide redistribution of H3K27me3 dominated by loss at promoters. A) CD1 mice were sacrificed at postnatal days 7, 14 and 60 after which cerebellar tissue was dissected out and processed to harvest histones, chromatin and RNA. B) (Left) Western blot of acid extracted histone from cerebellar tissue for H3K27me3 and total Histone H3 (n=3 biological replicates), (Right) Quantification of Western Blot, one-way ANOVA, * indicates p < 0.05. C) MA plot showing H3K27me3 peaks gained (Up) and lost (Down) throughout cerebellar maturation *in vivo* (n=3 biological replicates, differential enrichment calculated using DESeq2 package. Up and Down are loci with FDR < 0.05 and |L2FC| > 1). D) Percentage of differential H3K27me3 peaks between P7-P14, P14-P60 and P7-P60, annotated by genomic region (Annotation performed using ChIPseeker package, TSS +/- 3000 bp). E) (Left) Percentage of H3K27me3 peaks annotated by genomic region, (Right) Heatmap of H3K27me3 peaks centered around TSS +/- 5000 *bp*. F) ChIP-seq tracks of example ‘Down’ gene *Grin2c* (Upper) and ‘Up’ gene *Myc* (lower) (y-axes for ChIP-seq tracks are autoscaled according to data range in current view).

However, when we assessed the genomic location of H3K27me3 by ChIP-seq (Fig S1B) we observed a more complex pattern. We found local losses as well as gains in H3K27me3 enrichment at different sites between developmental stages (Fig 1C). This was true even when comparing P14 with P60 (Fig 1C), demonstrating that bidirectional changes in H3K27me3 distributions occur during postmitotic stages of neuronal maturation, well after the period of terminal cell fate commitment has ended (Fig S1C). We stratified differential H3K27me3 ChIP-seq peaks by their genomic location with respect to proximal gene promoters, gene bodies or intergenic regions (Fig 1D), and found a consistent overrepresentation of proximal gene promoters among developmentally demethylated peaks for all three comparisons. By contrast, H3K27me3 ChIP-seq peaks differentially gained between all three comparisons did not differ from the global distribution of H3K27me3, which was distributed across gene-body and distal intergenic regions as well as proximal promoters. The percentage of total H3K27me3-positive promoters gradually decreased by ~2-fold from P7 to P60, while that of H3K27me3-positive distal intergenic regions increased, and that of gene bodies stayed about the same over time (Fig 1E). Additionally, the total number of H3K27me3 peaks called centered within a 5 kb window around the TSS appeared to gradually decrease from P7 to P60 (Fig 1E). These data demonstrate that in CGNs H3K27me3 patterns are locally regulated in postmitotic neurons, suggesting a function for H3K27me3-dependent gene regulation in developing neurons that extends beyond the restriction of cell fate commitment.

### H3K27me3 Dynamics at Promoters During Cerebellar Maturation Inversely Correlate with Gene Expression

We filtered differential H3K27me3-peaks for those occurring at promoters because these regions can be directly mapped to their likely target gene for transcriptional regulation. We then asked whether peaks that were up- and downregulated over stages of CGN differentiation were found at genes that can be stratified according to cellular function. Examples of genes with promoter-associated peaks lost or gained from P7 to P60, respectively, were the genes encoding the mature NMDA-type glutamate receptor *Grin2c* which is strongly expressed in mature CGNs [11] and the cell-proliferation factor *Mycn*, which is expressed only in GNPs [36] (Fig 1F). This is consistent with the hypothesis that changes in the levels of promoter H3K27me3 inversely correlate with changes in gene expression over development.

To determine if the correlation of H3K27me3 demethylation with transcriptional maturation held more broadly, we first performed hierarchical clustering on all differential H3K27me3 ChIP-seq peaks and identified 3 major clusters with differing direction and timing of change (Fig 2A). The first cluster, which we termed ‘H3K27me3 Up,’ contained peaks that gained methylation as early as P14 and maintained methylation during CGN maturation. Gene Ontology (GO) analysis demonstrated that these peaks were enriched for genes associated with progenitor functional terms related to cell proliferation and morphogenesis. The next two clusters included demethylated peaks, that we termed ‘H3K27me3 Down (Fast)’ and ‘H3K27me3 Down (Slow)’ respectively, depending on whether they changed between P7 and P14 or between P14 and P60. Both clusters were enriched for neuronal GO terms. Because developmentally gained peaks showed a broader distribution than just promoters, we also performed the same type of filtering and hierarchical clustering for differential H3K27me3 peaks overlying gene bodies (Fig S2). This resulted in three clusters, two of which contained peaks that gained and lost methylation respectively at gene-bodies. Both these clusters were enriched for neuronal GO terms including early CGN-maturation genes. Together, these data show that bidirectional H3K27me3 dynamics are distinct across distinct genomic locations.

**Fig 2.**
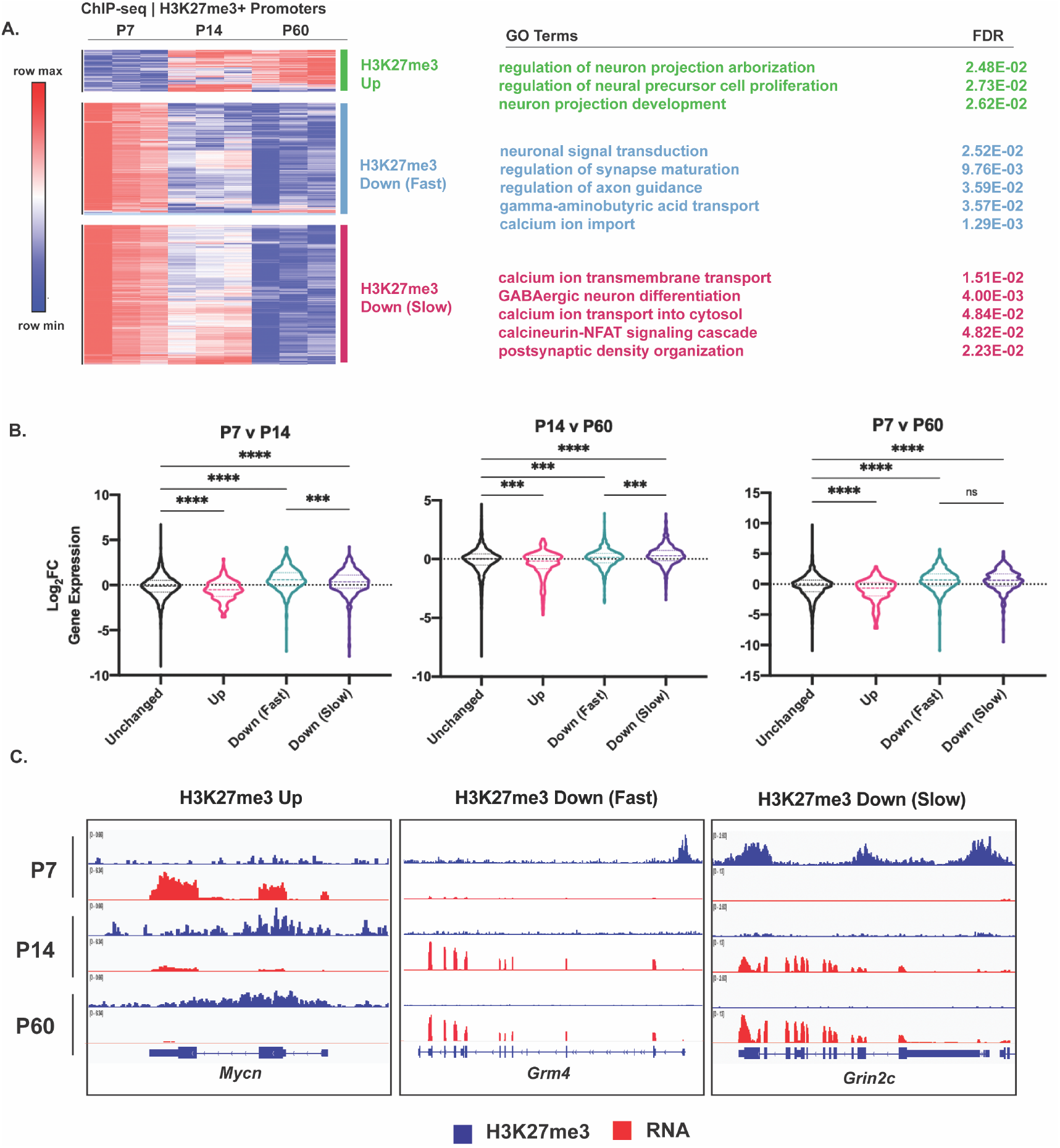
Differentially methylated promoters cluster by turnover kinetics and correlate with CGN maturation gene expression. A) (Left) Heatmap of hierarchically clustered, VST-transformed DESeq2-normalized counts of differential H3K27me3 promoter peaks from P7-P60, (Right) Corresponding Gene Ontology (GO) Terms associated with nearest gene and FDR. B) Violin plot showing the distribution of Log2FC of gene expression as a function of clustering performed in A, across developmental time (one-way ANOVA, * indicates p < 0.05). C) Example ChIP-seq tracks of genomic loci belonging to clusters described in A, at *Mycn, Grm4* and *Grin2c* loci and corresponding RNA-seq tracks (y-axes for ChIP-seq and RNA-seq tracks are autoscaled separately, according to data range in current view).

We then asked if H3K27me3 dynamics at gene promoters were associated with the expression of the corresponding RNA transcript across CGN maturation (Fig 2B, C). For this purpose, we compared our H3K27me3 ChIP-seq to a dataset of gene expression at the same time points from our previous study [27]. We found that ‘H3K27me3 Up’ peaks at gene promoters were significantly associated with a decrease in expression of the associated gene over the same period of developmental time and ‘H3K27me3 Down’ peaks were associated with an increase in gene expression. Among the H3K27me3 Down peaks, ‘Fast’ peaks were associated with a significantly greater magnitude of gene induction than ‘Slow’ peaks between P7 and P14, but ‘Slow’ peaks were associated with a significantly greater magnitude of gene induction than ‘Fast’ peaks between P14 and P60. There was no significant difference between ‘Fast’ and ‘Slow’ peaks from P7 to P60 indicating that it is the timing not the magnitude of differential developmental gene expression that defines the two clusters. These data show that the kinetics of H3K27me3 demethylation at gene promoters are associated with the timing of gene expression changes in postmitotic neurons, which allows for multiple postmitotic waves of gene expression (Fig. 2C).

### *Kdm6b-cKO* Impairs CGN Maturation via Promoter Hypermethylation

The developmental loss of H3K27me3 at a subset of gene promoters in CGNs could occur through changes in expression of the enzymes that maintain or remove these histone modifications. At the level of gene expression, RNA-seq counts of the H3K27 methyltransferases *Ezh1/2* and the H3K27 demethylases *Kdm6a/b* showed that although there was a developmental switch between the dominant reader and writer enzymes, the overall expression levels of each class of enzyme did not change over time (Fig S3). However, previous *in situs* from our lab demonstrated that *Kdm6b* expression is transiently induced in newborn CGNs in the inner EGL, and we further showed that shRNA-mediated knockdown of *Kdm6b* in cultured CGNs impairs the expression of a subset of developmentally induced genes [11]. Thus, we considered the possibility that KDM6B might mediate postmitotic loss of H3K27me3 at a subset of developmentally induced gene promoters in the developing cerebellum *in vivo* to promote their transcriptional induction. To test this hypothesis, we conditionally knocked out *Kdm6b* in cerebellar granule neuron precursor cells (GNPs) *in vivo* by crossing heterozygous transgenic *Atoh1-Cre* mice with *Kdm6b*^*fl*/fl^ mice. We then harvested cerebellar tissue from littermates that were either *Kdm6b*^*fl*/fl^ (WT) or *Atoh1-Cre; Kdm6b*^*fl*/fl^ (*Kdm6b-cKO*) at P14 as an intermediate developmental time point for CGN maturation [27] (Fig 3A). We processed this tissue to perform either RNA-seq (Fig S4A) or ChIP-seq for H3K27me3 (Fig S4B). We used DESeq2 to compute differentially enriched H3K27me3 ChIP peaks between WT and *Kdm6b-cKO* tissue at P14, and obtained 250 upregulated peaks, and 327 downregulated peaks (FDR < 0.05, |Log2FC| > 1) (Fig 3B). We then mapped these differential peaks to their genomic location (Fig 3C). We found peaks with elevated levels of H3K27me3 in *Kdm6b-cKO* mice to be significantly more likely to be within gene promoters compared with the overall distribution of peaks or the distribution of peaks that had reduced H3K27me3 in *Kdm6b-cKO* mice. We observed that the percentage of promoter associated H3K27me3 peaks decreased between WT and *Kdm6b-cKO* mice, while peaks in distal intergenic regions and gene-bodies increased (Fig S5C). The genes containing promoters with increased H3K27me3 in the *Kdm6b*-cKO were enriched for neuronal GO terms (Fig 3D), whereas those containing promoters with decreased H3K27me3 in the *Kdm6b*-cKO mice failed to pull any significant GO terms. Overall, these observations mirror our findings of H3K27 demethylation at the promoters of neuronal maturation genes and suggests that KDM6B mediates that process.

**Fig 3.**
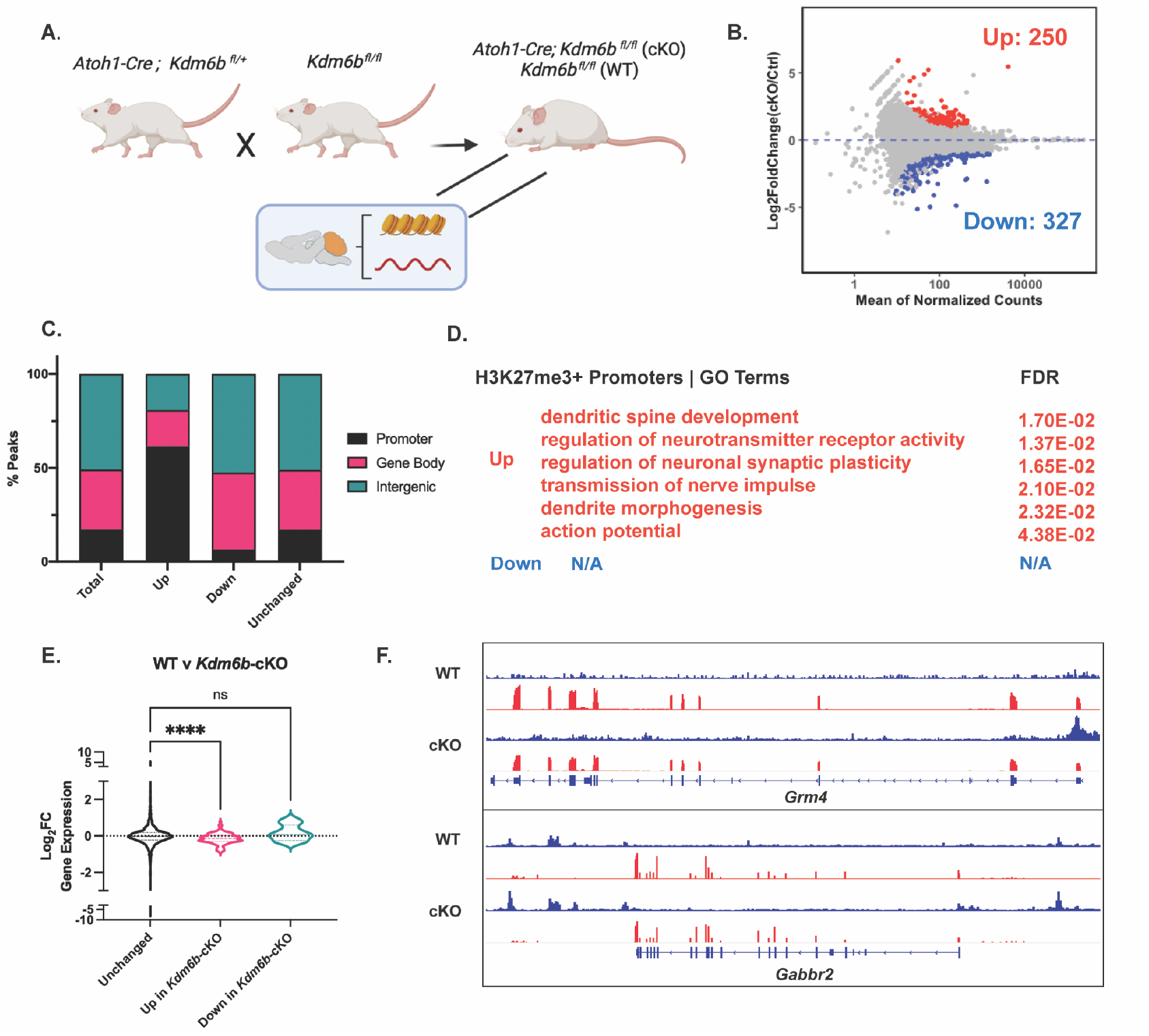
*Kdm6b* knockout in CGN precursors impairs CGN maturation via H3K27me3 hypermethylation of CGN maturation gene promoters. A) *Atoh1-Cre; Kdm6b*^*fl*/fl^ (cKO) and *Kdm6b*^*fl*/fl^ (WT) mice were generated by crossing *Atoh1-Cre; Kdm6b*^*fl*/+^ mice to *Kdm6b*^*fl*/fl^ mice and sacrificed at P14 for cerebellar tissue harvest and subsequent performing of H3K27me3 ChIP-seq (WT n=2, cKO n=3 biological replicates) and RNA-seq (n=2 biological replicates). B) MA plot showing differential enrichment of genome wide H3K27me3 peaks between WT and cKO mice (|Log2FC| >1, FDR < 0.05). C) Differentially methylated H3K27me3 between WT and cKO annotated by genomic location using ChIPseeker package, TSS +/- 3000 bp. D) GO Terms and FDR for genes nearest genomic promoters within H3K27me3 Up and Down (due to *Kdm6b*-cKO) peaks. E) Distribution of Log_2_Fold Change Expression (WT v cKO, computed using DESeq2) as a function of unchanged, H3K27me3 Up and H3K27me3 Down genomic promoters between WT and cKO (one-way ANOVA, * indicates p < 0.05). F) Representative genes containing H3K27me3 Up genomic promoter peaks *Grm4* and *Gabbr2* and corresponding RNA-seq tracks between WT and cKO mice (y-axes for ChIP-seq and RNA-seq tracks are autoscaled separately, according to data range in current view).

To determine the relationship between H3K27me3 levels at promoters and gene expression in the *Kdm6b*-cKO mice, we computed the correlation between *Kdm6b*-cKO associated differential H3K27me3 enrichment at promoter regions, with the expression of the associated gene in WT and *Kdm6b*-cKO mice. These data showed that the gain of H3K27me3 is associated with a significant loss of gene expression (Fig 3E) as exemplified by the synaptic genes *Grm4* and *Gabbr2* (Fig 3F). By contrast, promoters that lost H3K27me3 in *Kdm6b-cKO* mice showed no significant difference in gene expression (Fig 3E). These data suggest that developmental loss of H3K27me3 at the promoters of genes induced in CGNs during postmitotic maturation is facilitated by KDM6B, and that this loss promotes the developmental induction of gene expression.

### H3K27me3 Removal at CGN-Maturation Gene Promoters is Associated with H3K27ac Enrichment and Increased Chromatin Accessibility

Previously we showed CGN maturation to occur in concert with broad, genome-wide changes in H3K27ac [27] (Fig S5A). To ask whether H3K27me3 removal at promoters overlapped with the gain of H3K27ac, we computed the overlap between genes containing promoter peaks that were H3K27ac Up (| L2FC| > 0, FDR < 0.05) and H3K27me3 Down (Fig S5B). We observed that only a subset of genes that gained H3K27ac also lost H3K27me3 at P60. H3K27ac Up genes that were H3K27me3 Down (912 genes), or not H3K27me3 Down (4208 genes) were both highly enriched for neuronal GO terms (Fig S5C).

To determine what characterizes the subset of promoters that become demethylated we sought to ask what transcription factors might be found at these gene promoters compared to all H3K27me3-positive promoters. To conduct an unbiased screen, we first utilized the tool called Binding Analysis for Regulation of Transcription (BART) [37], which calculates enrichments of previously published ChIP-seq TF binding profiles within a defined set of genomic regions. We first filtered our ChIP-seq data for genes associated with the ‘H3K27me3-Down’ peak cluster described in Fig 2A, combining ‘Down (Fast)’, and ‘Down (Slow)’ clusters, and obtained predicted transcription factors (TF) whose binding significantly overlaps these gene promoters. We compared the enrichment statistics of TFs enriched in these clusters to those derived from all H3K27me3–promoter associated genes (Fig 4A). As expected, we found PRC2 components EZH2, JARID2, SUZ12 to be significantly enriched across both the H3K27me3-Down cluster and all H3K27me3-promoter associated genes. However, we found only the H3K27me3-Down cluster to be significantly enriched for the putative binding of transcription factors ZIC1 and ZIC2, which previous work from our group has shown to be a driver of CGN maturation [27]. Importantly, when we assessed the ‘H3K27me3-Down (Fast)’, and ‘H3K27me3-Down (Slow)’ clusters separately (Fig S6A), they were also enriched for the binding of ZIC1 and ZIC2, which is consistent with evidence [27] that these TFs function at multiple stages of CGN differentiation. By contrast, when we performed the same analysis for the ‘H3K27me3-Up’ cluster (Fig S6A), we found this set of promoters was significantly enriched for the binding of a distinct set of TFs involved in cell differentiation, namely SOX3, FOXF1, SHOX2, POU3F2 as well as histone deacetylation (HDAC1) [38].

**Fig 4.**
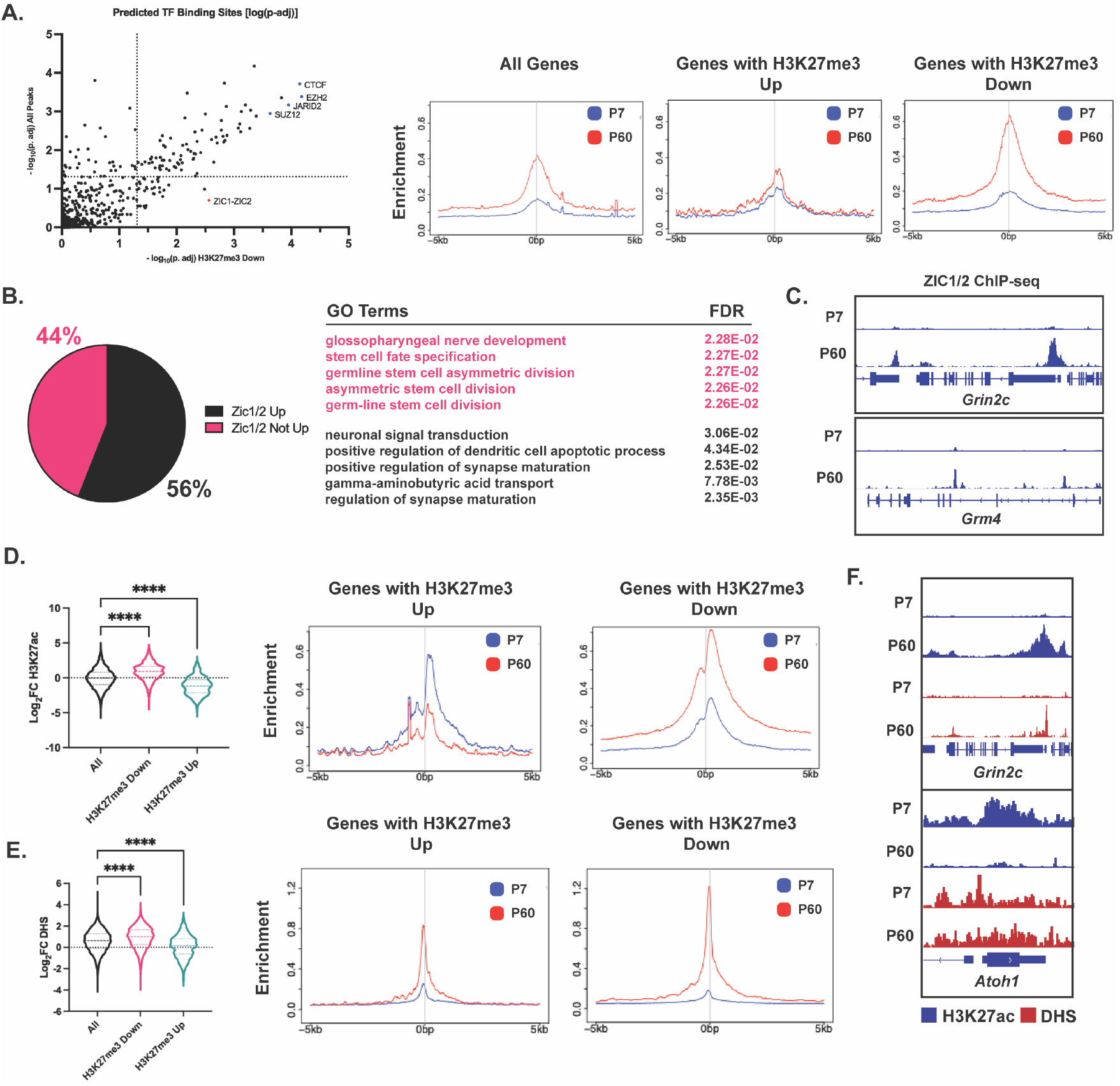
H3K27me3 removal is followed by gain of H3K27me3 and chromatin accessibility at gene promoters. A) (Left) Binding Analysis for Regulation of Transcription (BART) for genes near H3K27me3 Down peaks (x-axis) and near all H3K27me3 peaks (y-axis) plotted by −log_10_p-adj. (Right) Metagene plots for ZIC ChIP-seq signal at genes near all H3K27me3 peaks, genes with H3K27me3 Up and genes with H3K27me3 down. B) (Left) Percentage of genes near H3K27me3 Down promoter peaks that are also ZIC1/2 Up (Log2Fold Change > 1, P60/P7, ZIC1/2 ChIP-seq). (Right) GO Terms and FDR for genes nearest to H3K27me3 Down promoter peaks that are also ZIC1/2 Up, and not ZIC1/2 Up. C) ZIC ChIP-seq tracks at P7 and P60 at synaptic genes *Grm4* and *Grin2c* (y-axes for ChIP-seq are autoscaled separately, according to data range in current view). D) (Left) Distribution of Log2Fold Change of H3K27ac enrichment as a function of H3K27me3 Down and H3K27me3 Up described in Fig 2; and (Right) Metagene plots showing H3K27ac ChIP-seq signal at genes with H3K27me3 Up and H3K27me3 Down. E) (Left) Distribution of Log2Fold Change of DNase Hypersensitivity as a function of H3K27me3 Down and H3K27me3 Up described in Fig 2; and (Right) Metagene plots showing DNAse-seq signal at genes with H3K27me3 Up and H3K27me3 Down. F) H3K27ac ChIP-seq and DNAse-seq tracks at P7 and P60 at synaptic gene *Grin2c* (upper), and early gene *Atoh1* (lower) (y-axes for ChIP-seq are autoscaled separately, according to data range in current view).

To directly compare changes in ZIC1/2 TF binding to changes in H3K27me3 across CGN development, we re-analyzed our previously published ZIC1/2 ChIP-seq data [27] and measured ZIC1/2 binding at All Genes, Genes with H3K27me3 Up, and Genes with H3K27me3 Down (Fig 4A). We then identified gene promoters with increased ZIC binding at P60 compared with P7. We termed these genes ‘ZIC Up’ and first compared them to the H3K27me3-Down cluster. We found that 56% of H3K27me3-Down genes were also ZIC-Up, and that these genes were enriched for neuronal GO terms (Fig 4B). As a control, we compared ‘ZIC Up’ genes to H3K27me3-Up genes (Fig. S6A) and only found an overlap of only 11%. We repeated this analysis with the H3K27me3 ChIP-seq peaks that were differential at P14 between WT and *Kdm6b-cKO* (Fig. 3) and found a similar enrichment of putative ZIC1 and ZIC2 binding within gene promoters associated with H3K27me3 peaks ‘Up in *Kdm6b*-cKO’ (Fig S6B), but not ‘Down in *Kdm6b-cKO’* (Fig S6C). 72% of the ‘H3K27me3 Up in *Kdm6b*-cKO’ genes were also ‘ZIC Up’ (Fig S6B) and were enriched for neuronal GO terms. Only 9% of ‘H3K27me3 Down in *Kdm6b*-cKO’ were also ‘ZIC Up’ (Fig S6C). As examples of gene promoters that belong to both the H3K27me3-Down cluster, and H3K27me3 Up peaks due to *Kdm6b*-cKO, we show tracks for *Grin2c* and *Grm4*, that are strongly enriched for ZIC1/2 at their promoters at P60 (Fig 4C).

We have shown previously [27] that increased ZIC1/2 binding strongly correlates with the developmental enrichment of H3K27ac and increased chromatin accessibility during cerebellar maturation. Given the strong enrichment of ZIC1/2 binding within the H3K27me3-Down cluster during development, and H3K27me3 Up peaks due to *Kdm6b*-cKO, we wanted to ask whether H3K27me3 turnover during cerebellar maturation is associated with H3K27ac enrichment and changes in chromatin accessibility. As such we computed the distribution of Log2FC values from our previously published H3K27ac-ChIP-seq and DNAse Hypersensitivity data [27] comparing P7 and P60 cerebellum and filtered these distributions for those related to peaks belonging to our differential H3K27me3 promoter clusters. Indeed, H3K27me3 Down peaks at P60 were significantly associated with an increase in H3K27ac enrichment (Fig 4D) and DNAse hypersensitivity (Fig 4E). H3K27me3-Up peaks due to *Kdm6b*-cKO conversely were associated with genes that became enriched with H3K27ac, and DNAse hypersensitive regions at P60 (Fig S6D). As examples, we show H3K27ac ChIP-seq and DNAse-seq tracks for ‘H3K27me3 Down’ gene *Grin2c*, and ‘H3K27me3 Up’ gene *Atoh1*.Thus, these data provide further evidence that the postmitotic removal of H3K27me3 from a subset of gene promoters is important for the coordinated induction of gene expression that underlies neuronal maturation.

### Modulation of H3K27me3 Turnover Mediates CGN Maturation Gene-Regulation in Culture

To determine the functional consequences of modulating H3K27me3 at CGN-maturation gene promoters or gene bodies we used cultures of primary mouse CGNs as context for pharmacological manipulation of histone regulatory enzymes. When GNPs isolated from P6-8 mouse cerebellum are placed in culture they show changes in gene expression over the course of 7 days [27] (Fig 5A, S7A, B). We performed CUT&RUN-seq to measure genome-wide changes in H3K27me3 after 1, 3, 5 and 7 days *in vitro* (DIV) (Fig 5A). We found methylation to be gained quickly after cell cycle exit, with 1688 peaks significantly up between DIV1 and DIV3 CGNs, followed by 109 peaks between DIV3 and DIV5, and 2 peaks between DIV5 and DIV7 CGNs (Fig 5B). Like our observations *in vivo* (Fig 1, 2), we saw that demethylation occurred more gradually, with 477 peaks significantly down between DIV1 and DIV3 CGNs, followed by 227 and 25 peaks between DIV3 and DIV5, and DIV5 and DIV7 CGNs respectively. Again, consistent with our findings *in vivo*, we found demethylation to occur primarily at promoters (Fig 5C, Fig S7C), and gains in methylation to occur more uniformly, including at gene-bodies and distal intergenic regions. When we filtered for differential H3K27me3 peaks at gene promoters, they clustered in two groups that gain H3K27me3 or lose H3K27me3 at DIV7 (Fig 5D). Genes whose promoters were in the ‘Gain H3K27me3 at DIV7’ cluster were associated with cell fate specification GO terms as predicted by canonical PRC functions. Conversely, genes whose promoters were in the ‘Lose H3K27me3 at DIV7’ cluster were associated with functional neuronal GO terms. We observed mature synaptic genes such as *Grin2c* and *Grm4* to lose H3K27me3 from DIV1 to DIV7 (Fig 5E lower), and proliferation genes expressed in GNPs such as *Myc* and *Atoh1* to gain H3K27me3 from DIV1 to DIV7 in culture (Fig 5E upper). Finally, we stratified our previously published RNA-seq data between DIV0 and DIV7 CGNs by their respective methylation status (‘Gain H3K27me3 at DIV7’ or ‘Lose H3K27me3 at DIV7’ cluster) and calculated the distribution of Log2FC (Fig 5F).

**Fig 5.**
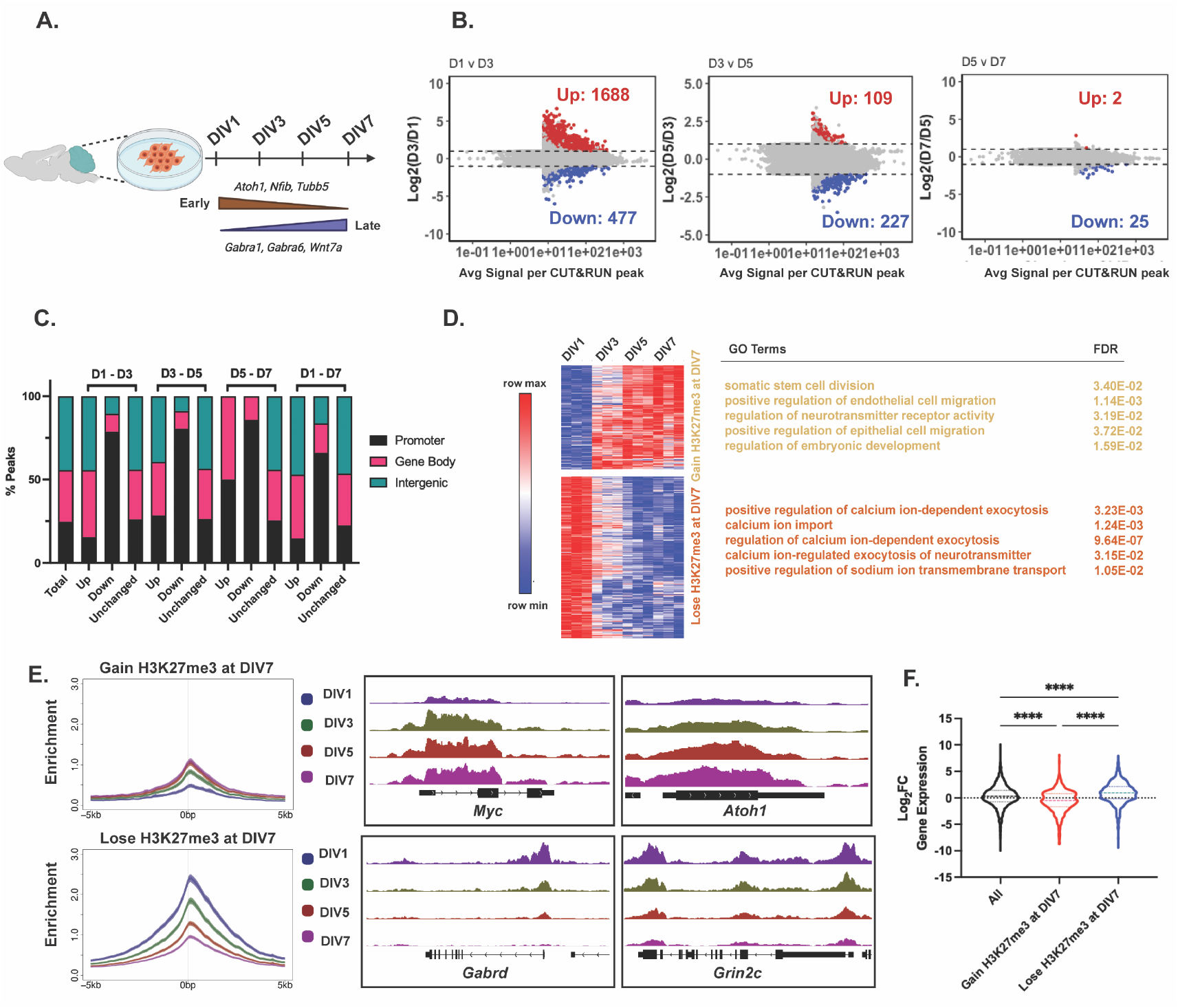
CGN Culture System refines temporal dynamics of H3K27me3 changes at gene promoters. A) Schematic describing CGN culture protocol over 7 days *in vitro* annotated with early and late CGN genes. B) Differentially methylated H3K27me3 CUT&RUN peaks annotated by genomic location, between DIV1-3, 3-5, 5-7 and 1-7 time points. C) Percentage of differential H3K27me3 CUT&RUN peaks between DIV1-3, DIV3-5, DIV5-7 and DIV1-7 annotated by genomic region (Annotation performed using ChIPseeker package, TSS +/- 3000 bp). D) (Left) Heatmap of hierarchical clustered VST-transformed DESeq2-normalized counts of differential H3K27me3 CUT&RUN promoter peaks from DIV1-7 (Right) Corresponding GO Terms and FDR associated with nearest gene. E) (Left) Metagene plots for H3K27me3 CUT&RUN signal at ‘Gain H3K27me3 at DIV7’ and ‘Lose H3K27me3 at DIV7’ clusters. (Right) CUT&RUN tracks for H3K27me3 for (top) early genes *Myc*, and *Atoh1* and (bottom) late genes *Gabrd* and *Grin2c* at DIV1, 3, 5 and 7 (y-axes for CUT&RUN tracks are autoscaled according to data range in current view). F) Violin plot showing the distribution of Log2FC of gene expression measured by RNA-seq between DIV0 and DIV7 CGNs as a function of clustering performed in E (one-way ANOVA, * indicates p < 0.05).

We found peaks associated with the ‘Gain H3K27me3 at DIV7’ cluster to have a significant reduction in gene expression over time in culture, and those associated with the ‘Lose H3K27me3 at DIV7’ cluster to have a significant gain in gene expression by DIV7 (Fig 5F, S7D, E). These data show that we can use this CGN culture system to explore mechanisms and causality with respect to the enzymes mediating H3K27me3 turnover and gene regulation during CGN maturation.

To determine the functional contribution of H3K27me3 regulation to CGN-maturation gene expression, we treated CGN cultures at DIV1 with small molecule inhibitors of either EZH2 or KDM6A/B, called GSK-126 and GSK-J4 respectively, and measured the consequences for cell viability, histone modification, and gene expression at DIV5. Because these drugs have not been widely used in primary neurons, we first performed a cell viability assay and measured the IC50 values of both drugs. We determined the IC50 values to be ~3.07 μM for GSK-J4 and ~7.32 μM for GSK-126, respectively (Fig S8A). Choosing concentrations below the measured IC50 values, we first measured global levels of H3K27me3 by western blot (Fig 6A), after treating with 0, 0.5, 1.0 and 2.0 μM of either drug. Treatment of CGNs with GSK-126 caused a dose-dependent loss of H3K27me3 validating the method. GSK-126 treatment also appeared to cause a global increase (Fig S8B) in H3K27ac, which is consistent with the idea that H3K27me3 and H3K27ac are mutually exclusive modifications that oppose each other’s functions [39]. By contrast, GSK-J4 treatment at these doses failed to cause any dose-dependent change in H3K27me3 detected by western or CUT&RUN (Fig. S8). Given that the IC_50_ was nearly 10-fold below the dose (25 μM) used to block the histone demethylase activity of the Kdm6s in cancer cells [40], we concluded that the therapeutic range is too small to permit its use in neurons.

**Fig 6.**
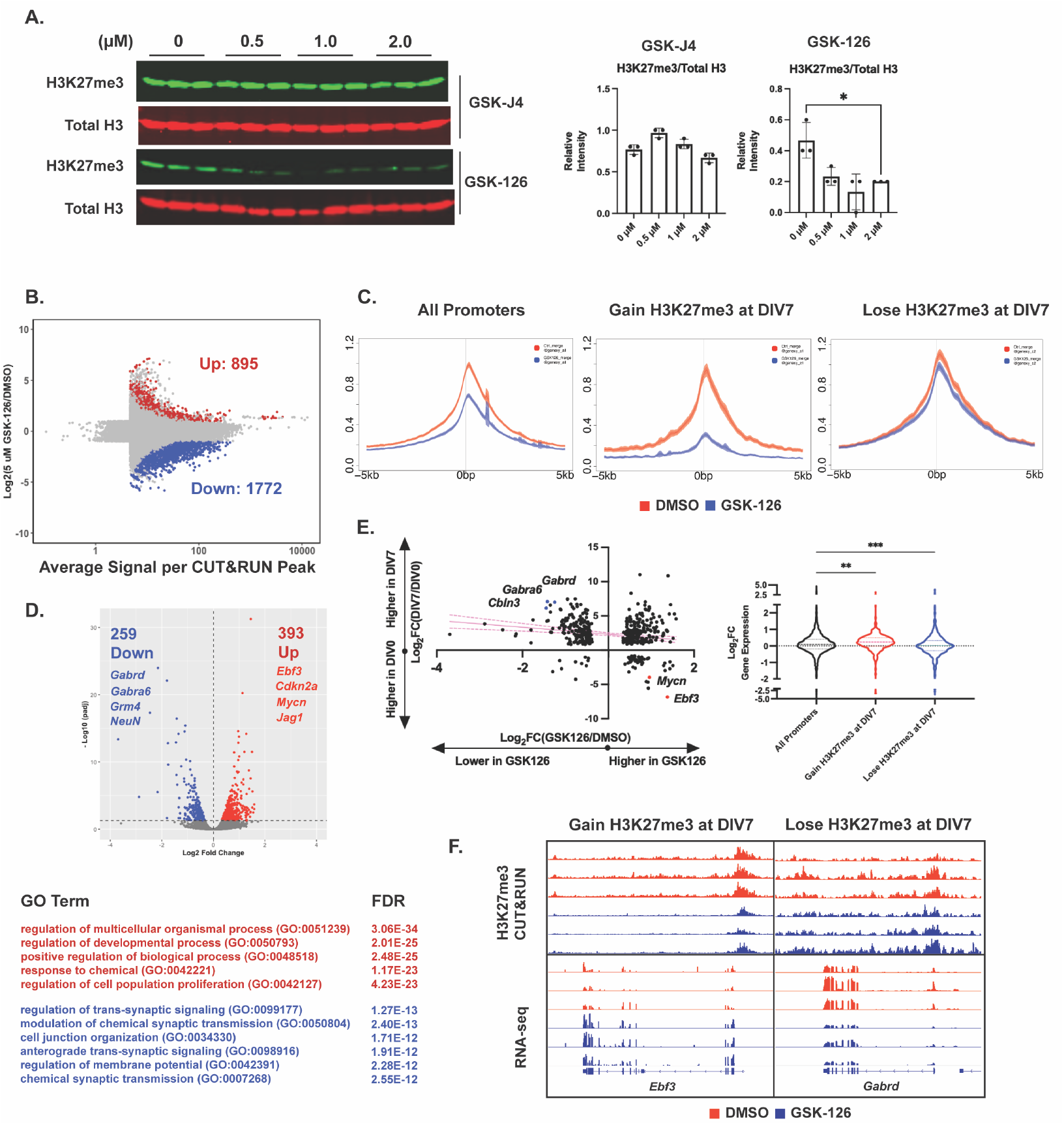
EZH2 catalytic activity temporally regulates CGN maturation in culture by depositing H3K27me3 at early CGN genes. A) (Left) Western blot of acid extracted histones from CGNs treated for H3K27me3 and total Histone H3 (n=3 biological replicates). CGNs were treated with GSK-J4 or GSK-126 at DIV1 and harvested at DIV5, (Right) Quantification of Western Blot. B) MA plot describing differential methylation due to GSK126 treatment in cultured CGNs (FDR < 0.05 and |L2FC| > 1) (n=3 biological replicates). C) Metagene plots for H3K27me3 peaks of DMSO and GSK-126 treated CGNs at genes near (Left) All H3K27me3 promoter peaks, (Center) genes that gain H3K27me3 at DIV7, and (Right) genes that lose H3K27me3 at DIV7, described in Fig 5E. D) (Upper) Volcano plot for differentially expressed genes measured by RNA-seq between DMSO and GSK-126 treated CGNs, (Lower) corresponding GO Terms and FDR for up- and downregulated genes. E) (Left) Relationship between gene expression changes during CGN maturation in culture and due to GSK-126 treatment. (Right) Violin plot showing the distribution of Log2FC of gene expression between DMSO and GSK-126 treated CGNs as a function of clustering performed in Fig 5D (one-way ANOVA, * indicates p < 0.05. F) (Upper) CUT and RUN tracks and (Lower) RNA-seq tracks for early genes *Ebf3* and synaptic gene *Gabrd*, for DMSO and GSK126 treated CGNs.

Our data showed that EZH2-inhibiting doses of GSK-126 were compatible with CGN viability, thus we then used CUT&RUN-seq for H3K27me3 on CGNs treated with 5 μM GSK-126 (Fig 6B, Fig S9A). Most of the regions that showed differential methylation following GSK-126 treatment had lower levels of H3K27me3 in GSK-126 treated neurons, consistent with the inhibition of this histone methyltransferase (Fig 6B). A much smaller number of peaks showed increases in H3K27me3, which may arise from secondary effects of EZH2 inhibition on other processes that impact H3K27me3 distributions such as EZH1 function [41] or histone turnover [42]. Regions that lost H3K27me3 after GSK-126 treatment showed a similar genomic distribution to the overall distribution of H3K27me3 peaks in CGNs (Fig 9B-C). Demethylated peaks were enriched for GO terms associated with several non-neuronal cell types, consistent with the general hypothesis that PRC functions in postmitotic cells at least in part to maintain the chosen cell fate [14] (Fig S9D).

To determine how EZH2 activity influences H3K27me3 distribution at early genes, we created metagene plots (Fig 6C) describing H3K27me3 coverage across all H3K27me3 peaks compared with the differential peaks from Fig 5D. We observed GSK-126 treatment to significantly reduce H3K27me3 coverage across the ‘Gain H3K27me3 at DIV7’ cluster but not the ‘Lose H3K27me3 at DIV7’ cluster or at all H3K27me3 peaks. To determine the effects of GSK-126 treatment on gene expression, we performed RNA-seq and identified 259 significantly downregulated and 393 significantly upregulated genes (Fig 6D, (upper)). Upregulated genes included *Mycn, Ebf3* and *Jag1* and were enriched for GO terms including ‘regulation of cell proliferation’, while downregulated genes included *Gabrd, Gabra6, Grm4* and were enriched for synaptic GO terms (Fig 6D, (lower)). We then computed gene expression profiles of CGNs maturing in culture (from DIV0 to DIV7) to CGNs treated with GSK-126 (Fig 6 left). We found genes repressed due to GSK-126 treatment to be strongly induced at DIV7 compared to DIV0. Conversely, we saw genes upregulated at DIV0 such as *Mycn* and *Ebf3* to be induced upon GSK-126 treatment. Furthermore, we computed changes to gene expression in response to GSK-126 treatment as a function of clusters described in Fig 5E – ‘All peaks’, ‘Gain H3K27me3 at DIV7’ and ‘Lose H3K27me3 at DIV7’ (Fig 6E right). We observed GSK-126 treatment to significantly upregulate the expression of genes belonging to the ‘Gain H3K27me3 at DIV7’ cluster, and significantly downregulate the expression of genes belonging to the ‘Lose H3K27me3 at DIV7’ cluster (Fig 6D, (upper right)). Visualizing H3K27me3 and RNA-seq tracks at early gene *Ebf3*, we observed GSK-126 treatment to completely block H3K27me3 and induce gene expression compared to DMSO treatment (Fig 6F upper). The late synaptic gene *Gabrd*, conversely, remained unchanged with respect to H3K27me3 enrichment, but was strongly downregulated in response to GSK-126 treatment (Fig 6F lower). These data show that EZH2 functions in newly postmitotic neurons to add H3K27me3 to genes that were not methylated at the time of cell fate commitment, and that this methylation helps to repress expression of progenitor genes [43] in neurons. By contrast, EZH2 is not required over this time course for the maintenance of H3K27me3 at genes that possessed the modification at the time of terminal differentiation, consistent with the slow turnover of methylated histones in postmitotic neurons. Why EZH2 catalytic activity early on during CGN maturation is critical for the induction of late genes is a point we discuss below.

## Discussion

In this study, we investigated the role of H3K27me3 turnover in postmitotic gene regulation, using the maturation of mouse CGNs as a model system. PRC2 activity has long been studied as a mechanism of early cell-fate specification in mammals through the deposition of H3K27me3 and silencing of fate defining genes during the differentiation of embryonic stem cells (ESCs) into other cell types including neural progenitor cells (NPCs) and early neurogenesis [5, 44, 45]. Our study demonstrates that dynamic changes in H3K27me3 continue in postmitotic neurons at specific sites in the genome to coordinate the temporal induction of gene expression programs that underlie functional maturation. Increases in H3K27me3 occurred rapidly after cell cycle exit, whereas promoter demethylation occurred gradually and these turnover kinetics strongly correlated with the kinetics of induced gene expression. Promoters and gene bodies that gained H3K27me3 after cell cycle exit were enriched for genes involved in proliferation and cell-fate specification such as *Atoh1* and *Myc*. By contrast, developmentally demethylated promoters included those for genes that confer mature function on synapses such as the NMDA receptor subunit *Grin2c* and the metabotropic glutamate receptor *Grm4*. This suggested to us that there are temporally coordinated mechanisms of enzymatic H3K27me3 turnover during CGN maturation, and that postmitotic H3K27me3 demethylation may be a key step for timing functional synaptic maturation. These data have important implications for understanding how ASD-associated mutations in the H3K27 lysine demethylase *KDM6B* may result in behavioral phenotypes.

Our *in vivo* cerebellar data show that developmental loss of H3K27me3 from gene promoters is facilitated by the H3K27me3 demethylase KDM6B, and further that conditional knockout of KDM6B in CGNs impaired the expression of genes that are induced as CGNs mature. *Kdm6b* expression is upregulated in newborn CGNs in the inner EGL [11], and we speculate that the temporal increase in expression of this enzyme may tip the balance of H3K27me3 regulation to favor histone demethylation at specific target sites. Our evidence that genes with increased H3K27me3 in the *Kdm6b-cKO* mice overlap those that show developmental loss of H3K27me3 is consistent with the possibility that KDM6B functions enzymatically to demethylate histones at these gene promoters. However, chromatin regulators including KDM6B have scaffolding functions as well as enzymatic functions [46–50], thus we attempted to use pharmacological inhibition of KDM6B to validate the role of its enzymatic activity in developing CGN gene regulation. The small molecule GSK-J4 is a highly selective inhibitor of the KDM6 family compared with other histone demethylases, and it was previously demonstrated in primary human macrophages to block LPS-mediated histone demethylation and induction of TNFa expression with an IC50 of 9 μM and no apparent toxicity up to 30 μM [40]. By contrast we found 50% cytotoxicity in CGNs at a GSK-J4 concentration of just 3.1 μM and no evidence by western blot or CUT&RUN for any global or local changes in H3K27me3 when the drug was applied at the non-toxic dose of 1 μM. Moving forward, as an alternative to pharmacological inhibition of KDM6B, genetic mutation of the JmjC domain of KDM6B may be a more effective means to test its functional requirement in neuronal H3K27me3 demethylation [51]. Indeed, a prior study showed that in a BAC transgene, enzymatically dead KDM6B was unable to rescue perinatal death in KDM6B knockout mice, which is consistent with a requirement for the enzymatic function of KDM6B in maturation of the respiratory circuit [9].

Our observation that H3K27me3 removal occurs at only a subset of genes that are developmentally induced during CGN maturation raises the question of how these promoters are marked for methylation and demethylation. Through an unbiased approach, we showed that promoters that lose H3K27me3 during CGN maturation are strongly enriched for binding of the transcription factors ZIC1/2; thus, one possibility is that an interaction between KDM6B and the ZICs selects these genes for induction in maturing neurons. ZIC1/2 are strong drivers of CGN maturation [27], and mutation of these genes is associated with cerebellar disorders, showing their importance in CGN development [52, 53]. ZIC1 and 2 are zinc-finger binding proteins that exhibit dynamic DNA binding properties during cerebellar maturation, and they regulate the expression of both early and late CGN maturation genes [27]. Given the strong association of H3K27ac enrichment at these promoters, however, it is likely that ZIC1/2 binding is a consequence and not a cause of H3K27-demethylation at promoters. It would therefore be essential to measure the genome-wide enrichment patterns of KDM6B during CGN maturation, in order to identify its binding partners and direct targets. Existing literature investigating KDM6B dynamics across the epigenome is sparse owing to the lack of well validated, commercially available KDM6B antibodies. Nevertheless, KDM6B has been shown to localize to promoters and enhancers of neurogenic genes in NSCs [54, 55], and more recently to induce muscle regeneration gene *Has2*, by removing H3K27me3 from its promoter, to initiate muscle repair in response to injury [51]. KDM6B targeting to these specific loci might be facilitated through interactions with TFs [55, 56], and chromatin remodeling complexes [57]. For example, KDM6B has been shown to strongly associate with the ATP-dependent chromatin remodelers mSWI/SNF or BAF complex, by interacting with its subunits which in turn potentiate its activity [58]. This interaction has been shown to play a role in neuronal development [58, 59].

Interestingly, we find that maturation of CGN transcriptional programs not only depends on the loss of H3K27me3 at genes that turn on late, but also requires the addition of H3K27me3 at regions that gain this mark in newly postmitotic neurons. Using CUT&RUN in cultured CGNs, we discovered that the rapid increases in H3K27me3 that accumulate in newly postmitotic neurons are highly sensitive to inhibition of EZH2 with GSK-126. Failure to gain H3K27me3 at these sites in differentiating neurons not only prevented the developmental downregulation of CGN progenitor markers like *Mycn* and *Ebf3* but also impaired the developmental induction of late genes like *Gabrd*, *Grm4* and *Grin2c*. The impact of GSK-126 on the mature CGN transcriptional program was not due to changes in H3K27me3 regulation at the late genes themselves. This shows that the postmitotic addition of H3K27me3 at specific genomic sites is crucial in regulating the induction of genes that turn on well after neuronal cell fate is determined.

However, these data raise the question of how PRC2 complex function might indirectly regulate the activation of late gene expression programs over a period of days. One possibility is that the persistent expression of early TFs like MYCN and EBF3 interferes with the expression or function of the TFs that activate late gene promoters even once these promoters lose H3K27me3. However, another possibility may be via the effects of H3K27me3 on chromatin architecture. Though H3K27me3 is an acutely repressive mark, PRC2 dependent deposition of H3K27me3 has been shown in some cases to contribute to physical interactions between enhancers and their target genes by anchoring chromatin loops [60, 61]. These loops may then poise genes for enhancer activation later upon receiving a developmental stimulus. The gain of H3K27me3 in early differentiating CGNs could serve a similar function to establish such enhancer contacts with the promoters of late genes such that when KDM6B demethylates these promoters, the enhancer can permit activation. In the future, investigating dynamics in higher-order chromatin structure through low-input techniques such as HiCAR [62] might help inform how H3K27me3 dynamics contribute to gene regulation in postmitotic neurons.

