## Supplementary Figures for "Bidirectional regulation of postmitotic H3K27me3 distributions underlie cerebellar granule neuron maturation dynamics"

**Figure S1. H3K27me3 ChIP-seq and RNA-seq profile of the Developing Cerebellum *in vivo***
A) (upper) Western blot of acid extracted histone from cerebellar tissue for H3K27ac and total Histone H3 (n=3 biological replicates), (lower) Quantification of Western Blot Quantification of Western Blot, one-way ANOVA, * indicates $p < 0.05$. B) Sample to Sample Distance for H3K27me3 ChIP-seq peaks for P7, P14 and P60 tissues and their 3 biological replicates. C) (Left) Heatmap of hierarchical clustered VST-transformed DESeq2-normalized RNA-seq counts for significantly different genes from P7-P60, (Right) Corresponding Gene Ontology (GO) Terms and FDR associated with each cluster.

**Figure S2. Differential H3K27me3 Enrichment at Gene Body Regions and Corresponding GO Terms**
**A)** (Left) Heatmap of hierarchical clustered VST-transformed DESeq2-normalized counts of differential H3K27me3 gene-body peaks from P7-P60, (Right) Corresponding Gene Ontology (GO) Terms and FDR associated with nearest gene.

**Figure S3. Expression of H3K27me3 Writers and Erasers *in vivo***
A) VST-transformed DESeq2-normalized RNA-seq counts for H3K27me3 Writers *Ezh1*/2 and B) *Kdm6a*/*b*.

**Figure S4. *Kdm6b*-cKO in *Atoh1-Cre*+ GNPs Impairs CGN Maturation *in vivo***
A) Heatmap and hierarchical clustered VST-transformed DESeq2-normalized RNA-seq counts of differential genes between WT and *Kdm6b*-cKO mice (left) and corresponding GO terms and FDR for those clusters (right) (n=2 biological replicates). B) Sample to sample distance for H3K27me3-peaks derived from WT and *Kdm6b*-cKO tissue and their biological replicates (n=3 biological replicates). C) (left) Percentage of H3K27me3 peaks annotated by genomic location – promoter, gene body or distal intergenic region. (right) Heatmap showing H3K27me3 peak coverage between WT and cKO tissue with a window of TSS +/- 5 kb.

**Figure S5. Most H3K27me3 Down Peaks Gain H3K27ac at P60**
A) MA plot describing differential H3K27ac enrichment between P7 and P70 cerebellum. $|L2FC > 0, FDR < 0.05|$. Data derived from [11]. B) Venn Diagram showing overlap between genes under 'H3K27ac Up' promoter peaks and 'H3K27me3 Down' promoter peaks. C) GO Terms and FDR for 'H3K27ac Up $\cup$ H3K27me3 Down' genes and 'H3K27ac Up' genes.

**Figure S6. H3K27me3 Turnover at CGN-Maturation Gene Promoters is Associated with Gain of Zic Binding**
A) (upper) BART analysis for genes nearest to peaks within H3K27me3 Up, H3K27me3 Down (Fast) and H3K27me3 Down (Slow) clusters described in Fig 2A versus genes nearest to all H3K27me3 promoter peaks plotted as –log10(p-adj). (lower) Overlap between genes nearest to peaks within H3K27me3 Up, H3K27me3 Down (Fast) and

H3K27me3 Down (Slow) clusters and ZIC1/2 Up promoter peaks. B) B) (Left) BART analysis comparing genes H3K27me3 Up peaks due to *Kdm6b*-cKO described in Fig 3. (Center) Percentage of genes near H3K27me3 Up peaks due to *Kdm6b*-cKO that are also ZIC1/2 Up ($Log_2$Fold Change > 1, P60/P7, ZIC1/2 ChIP-seq). (Right) GO Terms and FDR for genes nearest to H3K27me3 Up peaks due to *Kdm6b*-cKO that are also ZIC1/2 Up, and not ZIC1/2 Up. C) BART analysis for genes nearest to H3K27me3-Down peaks due to Kdm6b-cKO vs all H3K27me3 promoter peaks plotted as –log10(p-adj), (right) overlap between genes nearest to ZIC1/2 Up promoter peaks. D) Distribution of $Log_2$Fold Change of H3K27ac enrichment and DNAse Hypersensitivity as a function of 'Up in *Kdm6b*-cKO' and 'Down in *Kdm6b*-cKO' described in Fig 3 (one-way ANOVA, * indicates $p < 0.05$).

**Figure S7. CUT&RUN-seq Captures Genome-Wide Changes in H3K27me3 in Cultured CGNs**
A) Sample to sample distance for H3K27me3 CUT&RUN peaks from CGNs at DIV1, 3, 5 and 7 and their biological replicates. B) RT-qPCR for (top) early genes *Atoh1, Nfib* and *Tubb5* and (bottom) late genes *Gabra1*, *Wnt7a* and *Gabra6* (n=2-3 biological replicates). C) Percentage of H3K27me3 peaks annotated by genomic location – promoter, gene body or distal intergenic region. D) VST-transformed DESeq2-normalized RNA-seq counts for early genes *Atoh1*, *Nfib* and *Tubb5* (left) and late genes *Grin2c*, *Grm4* and Wnt7a (right). E) VST-transformed DESeq2-normalized RNA-seq counts for H3K27me3 Writers *Ezh1*/2 (left) and *Kdm6a/b* (right).

**Figure S8**. **GSK-J4 Treatment at 1 µM does not Affect Genome Wide Levels of H3K27me3, but Influences Expression of Late CGN-genes**
A) Cell Viability Assay for CGNs treated with GSK-126 or GSK-J4 at DIV1 and measured at DIV5 (n=2 biological replicates). B) Western blot of acid extracted histones from CGNs treated for H3K27ac and total Histone H3 (n=3 biological replicates). C) Percentage of H3K27me3 peaks annotated by genomic location – promoter, gene body or distal intergenic region. D) Sample to sample distance for H3K27me3 CUT&RUN peaks for DMSO and GSK-J4 treated samples and their biological replicates. E) MA Plot showing differential peak enrichment between DMSO and GSK-J4 treated CGNs. F) RT-qPCR for early genes *Tubb5*, *Nfib* and *Myc* (left) and late genes *Olfm3*, *Vglut2* and *Grin2c* (right) for CGNs treated with DMSO or GSK-J4 (n=3 biological replicates).

**Figure S9. EZH2 Inhibition Redistributes H3K27me3 Across CGN Genome**
A) Sample to sample distance for H3K27me3 CUT and RUN peaks for DMSO and GSK-126 treated CGNs in cultured and their 3 biological replicates. B) (left) H3K27me3 peaks annotated by genomic location – promoter, gene body and distal intergenic regions. (right) Heatmap showing H3K27me3 peak coverage for CGNs treated with DMSO or GSK-126 within a window of TSS +/- 5 kb (left), Percentage of H3K27me3 peaks annotated by genomic location – promoter, gene body or distal intergenic region (right). C) Percentage of differential H3K27me3 peaks between DMSO and GSK-126 treated CGNs, annotated by genomic region. D) GO Terms and FDR associated with genes nearest to peaks described as 'Up' or 'Down' in Figure 6.

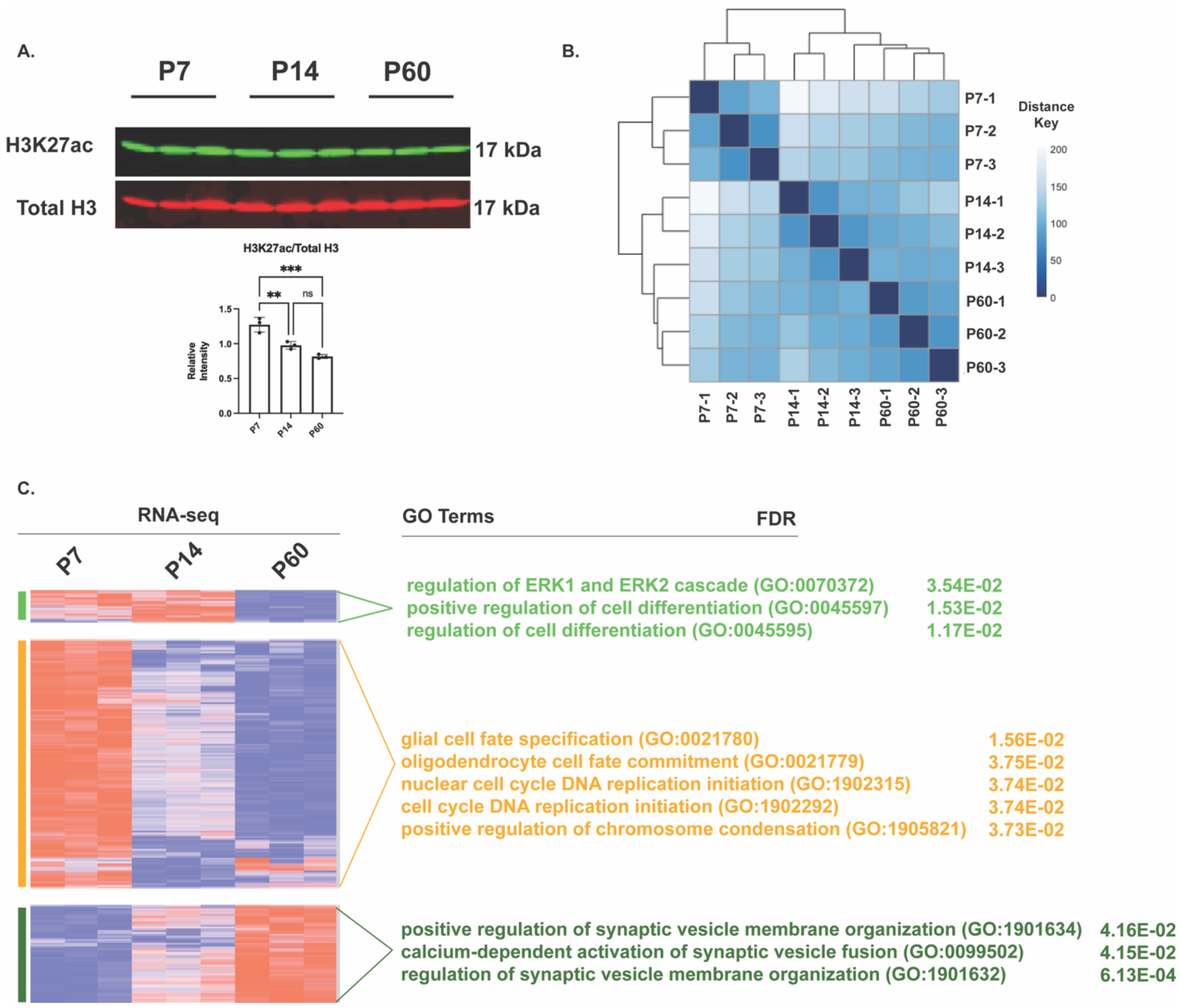


**Figure S1. H3K27me3 ChIP-seq and RNA-seq profile of the Developing Cerebellum *in vivo***

A) (upper) Western blot of acid extracted histone from cerebellar tissue for H3K27ac and total Histone H3 (n=3 biological replicates), (lower) Quantification of Western Blot Quantification of Western Blot, one-way ANOVA, * indicates $p < 0.05$. B) Sample to Sample Distance for H3K27me3 ChIP-seq peaks for P7, P14 and P60 tissues and their 3 biological replicates. C) (Left) Heatmap of hierarchical clustered VST-transformed DESeq2-normalized RNA-seq counts for significantly different genes from P7-P60, (Right) Corresponding Gene Ontology (GO) Terms and FDR associated with each cluster.

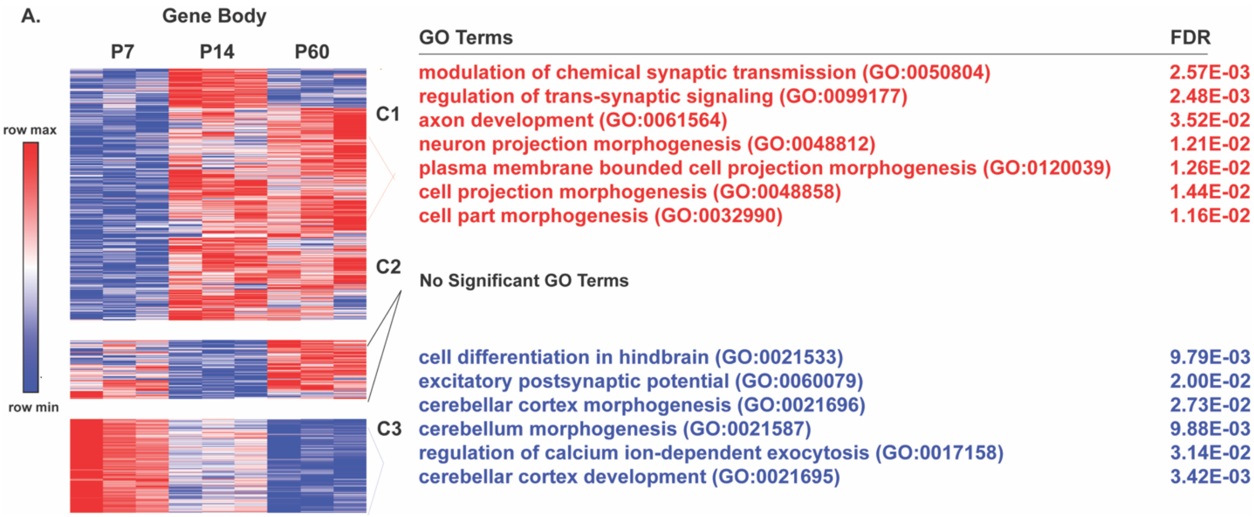


**Figure S2. Differential H3K27me3 Enrichment at Gene Body Regions and Corresponding GO Terms**

**A)** (Left) Heatmap of hierarchical clustered VST-transformed DESeq2-normalized counts of differential H3K27me3 gene-body peaks from P7-P60, (Right) Corresponding Gene Ontology (GO) Terms and FDR associated with nearest gene.

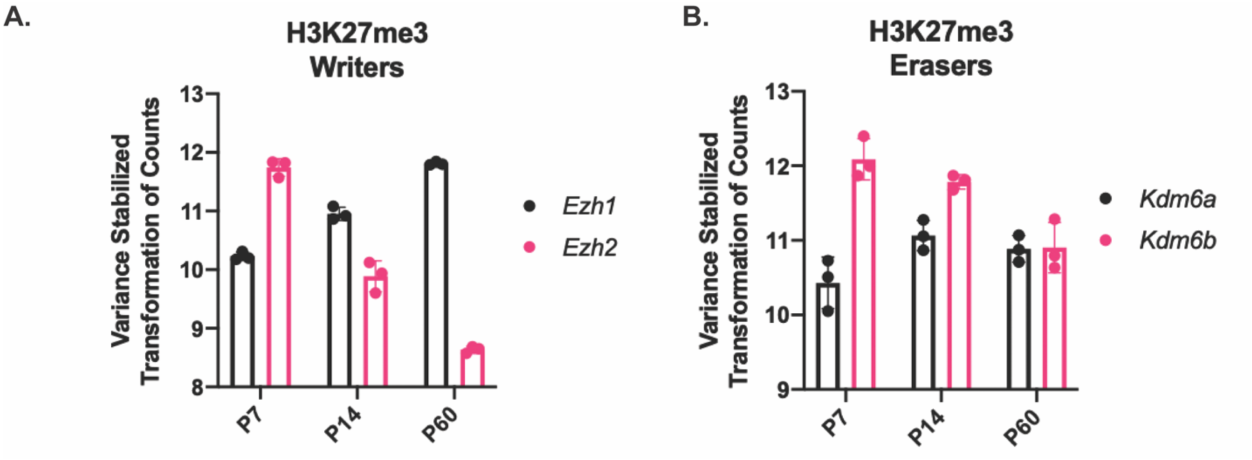


**Figure S3. Expression of H3K27me3 Writers and Erasers *in vivo***
A) VST-transformed DESeq2-normalized RNA-seq counts for H3K27me3 Writers *Ezh1*/2 and B) *Kdm6a*/*b*.

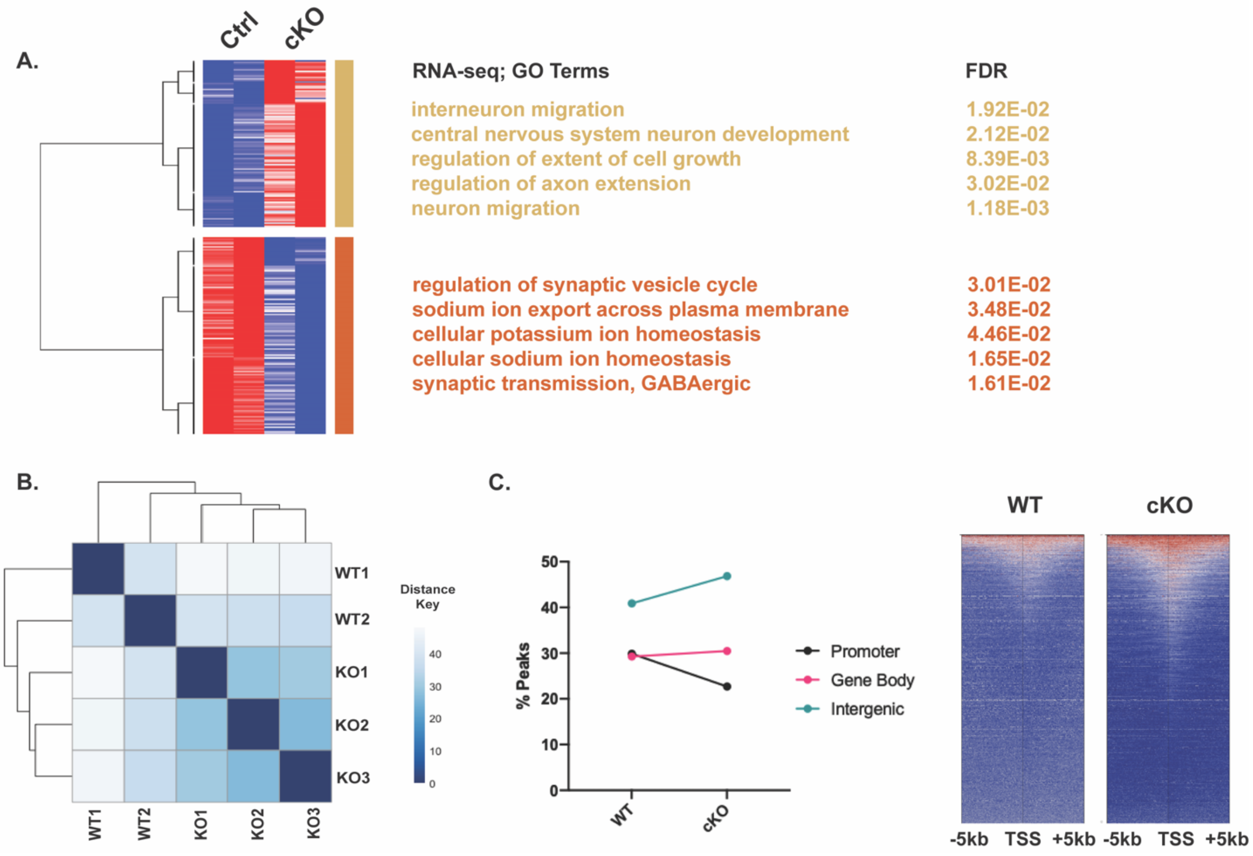


**Figure S4. *Kdm6b*-cKO in *Atoh1-Cre*+ GNPs Impairs CGN Maturation *in vivo***
A) Heatmap and hierarchical clustered VST-transformed DESeq2-normalized RNA-seq counts of differential genes between WT and *Kdm6b*-cKO mice (left) and corresponding GO terms and FDR for those clusters (right) (n=2 biological replicates). B) Sample to sample distance for H3K27me3-peaks derived from WT and *Kdm6b*-cKO tissue and their biological replicates (n=3 biological replicates). C) (left) Percentage of H3K27me3 peaks annotated by genomic location – promoter, gene body or distal intergenic region. (right) Heatmap showing H3K27me3 peak coverage between WT and cKO tissue with a window of TSS +/- 5 kb.

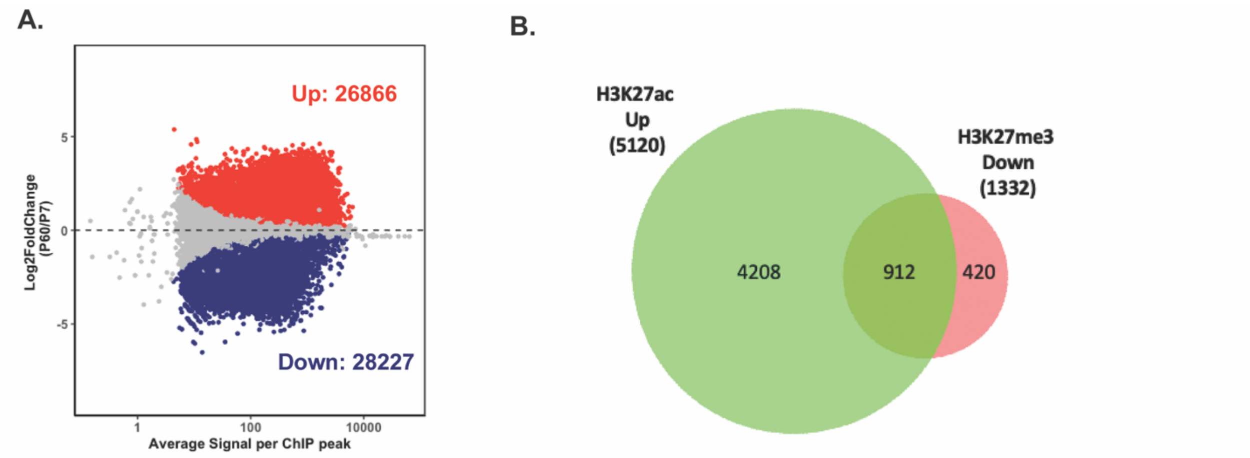


C.

| GO Terms: (912 genes) | FDR | GO Terms: (4208 genes) | FDR |
|---|---|---|---|
| neuronal signal transduction (GO:0023041) | 4.70E-02 | synaptic vesicle docking (GO:0016081) | 2.49E-02 |
| reelin-mediated signaling pathway (GO:0038026) | 2.51E-02 | synaptic vesicle priming (GO:0016082) | 4.78E-02 |
| regulation of long-term synaptic depression (GO:1900452) | 3.09E-04 | regulation of synaptic vesicle endocytosis (GO:1900242) | 3.64E-02 |
| regulation of synapse maturation (GO:0090128) | 5.49E-03 | regulation of synaptic vesicle recycling (GO:1903421) | 1.56E-02 |
| long-term synaptic potentiation (GO:0060291) | 1.00E-07 | synaptic membrane adhesion (GO:0099560) | 3.33E-02 |

**Figure S5**. **Most H3K27me3 Down Peaks Gain H3K27ac at P60**
A) MA plot describing differential H3K27ac enrichment between P7 and P70 cerebellum. |L2FC > 0, FDR < 0.05|. Data derived from [11]. B) Venn Diagram showing overlap between genes under 'H3K27ac Up' promoter peaks and 'H3K27me3 Down' promoter peaks. C) GO Terms and FDR for 'H3K27ac Up ∪ H3K27me3 Down' genes and 'H3K27ac Up' genes.

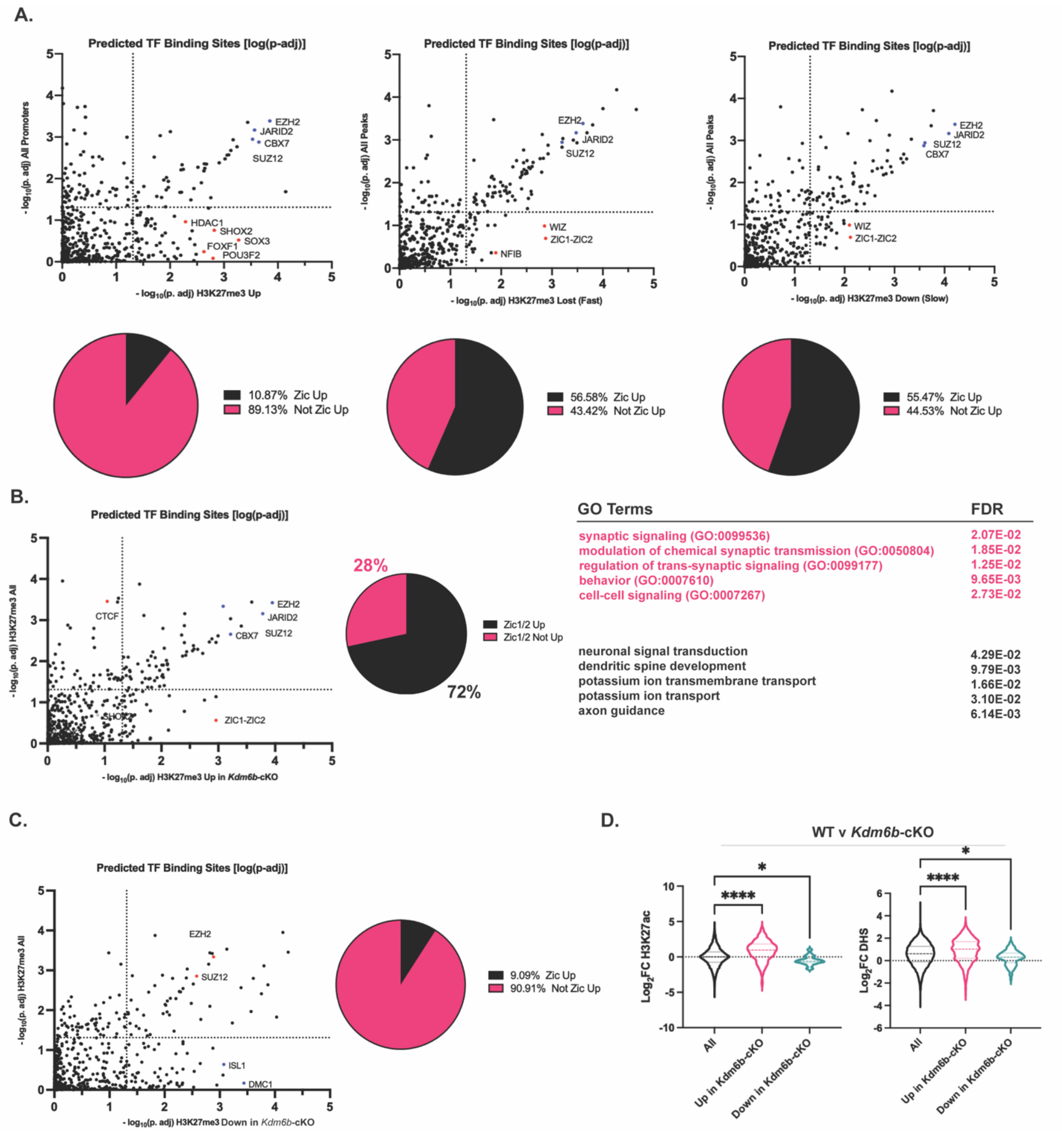


**Figure S6. H3K27me3 Turnover at CGN-Maturation Gene Promoters is Associated with Gain of Zic Binding**

A) (upper) BART analysis for genes nearest to peaks within H3K27me3 Up, H3K27me3 Down (Fast) and H3K27me3 Down (Slow) clusters described in Fig 2A versus genes nearest to all H3K27me3 promoter peaks plotted as –log10(p-adj). (lower) Overlap between genes nearest to peaks within H3K27me3 Up, H3K27me3 Down (Fast) and H3K27me3 Down (Slow) clusters and ZIC1/2 Up promoter peaks. B) B) (Left) BART analysis comparing genes H3K27me3 Up peaks due to *Kdm6b*-cKO described in Fig 3.

(Center) Percentage of genes near H3K27me3 Up peaks due to *Kdm6b*-cKO that are also ZIC1/2 Up ($Log_2$Fold Change > 1, P60/P7, ZIC1/2 ChIP-seq). (Right) GO Terms and FDR for genes nearest to H3K27me3 Up peaks due to *Kdm6b*-cKO that are also ZIC1/2 Up, and not ZIC1/2 Up. C) BART analysis for genes nearest to H3K27me3-Down peaks due to Kdm6b-cKO vs all H3K27me3 promoter peaks plotted as –log10(p-adj), (right) overlap between genes nearest to ZIC1/2 Up promoter peaks. D) Distribution of $Log_2$Fold Change of H3K27ac enrichment and DNAse Hypersensitivity as a function of 'Up in *Kdm6b*-cKO' and 'Down in *Kdm6b*-cKO' described in Fig 3 (one-way ANOVA, * indicates $p < 0.05$).

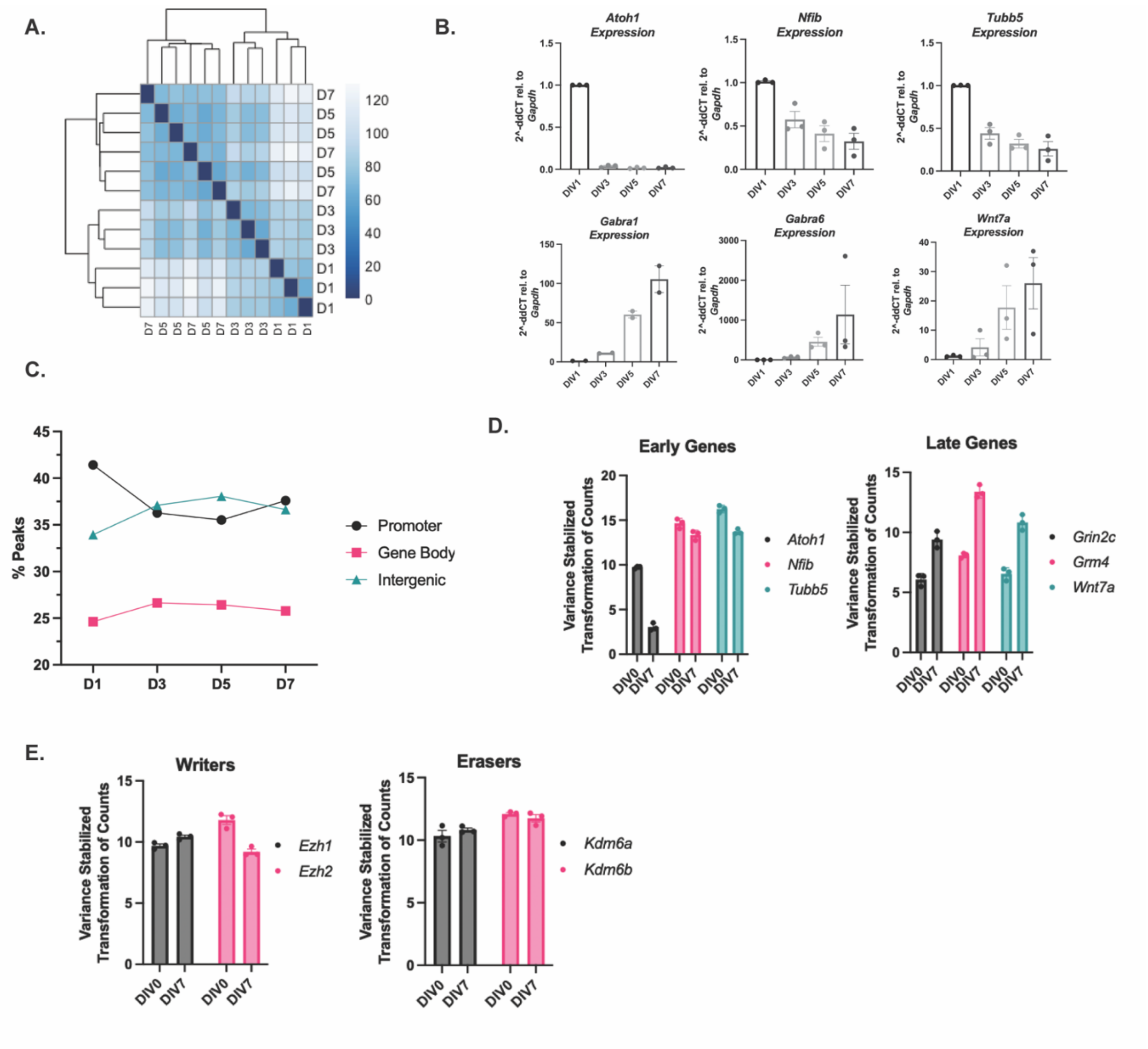


**Figure S7. CUT&RUN-seq Captures Genome-Wide Changes in H3K27me3 in Cultured CGNs**

A) Sample to sample distance for H3K27me3 CUT&RUN peaks from CGNs at DIV1, 3, 5 and 7 and their biological replicates. B) RT-qPCR for (top) early genes *Atoh1, Nfib* and *Tubb5* and (bottom) late genes *Gabra1*, *Wnt7a* and *Gabra6* (n=2-3 biological replicates). C) Percentage of H3K27me3 peaks annotated by genomic location – promoter, gene body or distal intergenic region. D) VST-transformed DESeq2-normalized RNA-seq counts for early genes *Atoh1*, *Nfib* and *Tubb5* (left) and late genes *Grin2c*, *Grm4* and Wnt7a (right). E) VST-transformed DESeq2-normalized RNA-seq counts for H3K27me3 Writers *Ezh1*/2 (left) and *Kdm6a/b* (right).

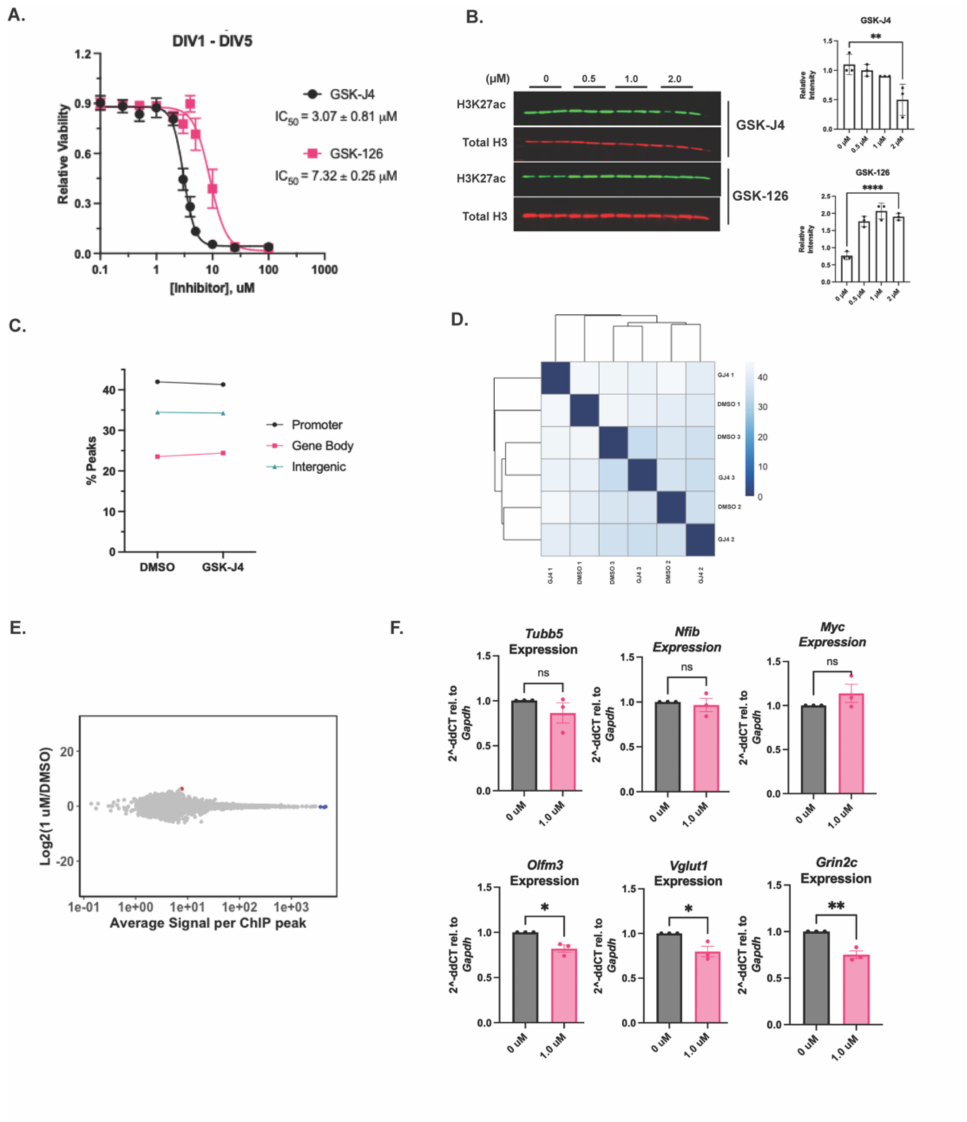


**Figure S8**. **GSK-J4 Treatment at 1 µM does not Affect Genome Wide Levels of H3K27me3, but Influences Expression of Late CGN-genes**

A) Cell Viability Assay for CGNs treated with GSK-126 or GSK-J4 at DIV1 and measured at DIV5 (n=2 biological replicates). B) Western blot of acid extracted histones from CGNs treated for H3K27ac and total Histone H3 (n=3 biological replicates). C) Percentage of H3K27me3 peaks annotated by genomic location – promoter, gene body or distal intergenic region. D) Sample to sample distance for H3K27me3 CUT&RUN peaks for DMSO and GSK-J4 treated samples and their biological replicates. E) MA Plot showing differential peak enrichment between DMSO and GSK-J4 treated CGNs. F) RT-qPCR for early genes *Tubb5*, *Nfib* and *Myc* (left) and late genes *Olfm3*, *Vglut2* and *Grin2c* (right) for CGNs treated with DMSO or GSK-J4 (n=3 biological replicates).

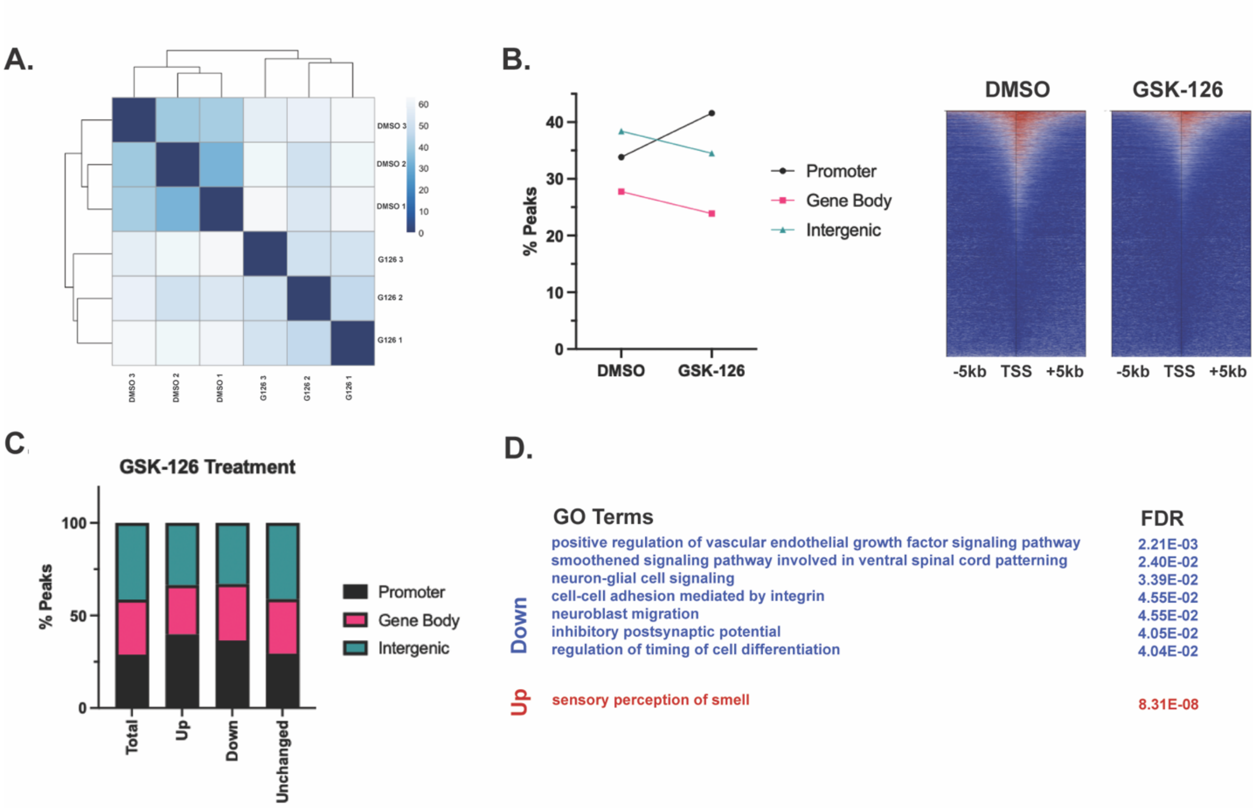


**Figure S9. EZH2 Inhibition Redistributes H3K27me3 Across CGN Genome**
A) Sample to sample distance for H3K27me3 CUT and RUN peaks for DMSO and GSK-126 treated CGNs in cultured and their 3 biological replicates. B) (left) H3K27me3 peaks annotated by genomic location – promoter, gene body and distal intergenic regions. (right) Heatmap showing H3K27me3 peak coverage for CGNs treated with DMSO or GSK-126 within a window of TSS +/- 5 kb (left), Percentage of H3K27me3 peaks annotated by genomic location – promoter, gene body or distal intergenic region (right). C) Percentage of differential H3K27me3 peaks between DMSO and GSK-126 treated CGNs, annotated by genomic region. D) GO Terms and FDR associated with genes nearest to peaks described as 'Up' or 'Down' in Figure 6.
