## Supplementary material for "Bidirectional regulation of postmitotic H3K27me3 distributions underlie cerebellar granule neuron maturation dynamics": Table S1

Table S1. PCR and qPCR Primers

| **Primer** | **Organism** | **Assay** | **Sequence (5’ – 3’)** |
| --- | --- | --- | --- |
| *Gapdh* F | Mouse | qPCR | CATGGCCTTCCGTGTTCCT |
| *Gapdh* R | Mouse | qPCR | TGATGTCATCATACTTGGCAGGTT |
| *Grin2c* F | Mouse | qPCR | TGTGGGCCTTCTTCGCTGTCATCT |
| *Grin2c* R | Mouse | qPCR | TGCCATTAGGTACCGTGCCAAAAC |
| *Wnt7a* F | Mouse | qPCR | GCTGCCTGGGCCACCTCTTTCTCA |
| *Wnt7a* R | Mouse | qPCR | CCCGGTGGTACTGGCCTTGCTTCT |
| *Grm4* F | Mouse | qPCR | CTTCCTTAGCCAGGGTCTCC |
| *Grm4* R | Mouse | qPCR | CATCCCTTCGGACACAGTTT |
| *Tubb5* 5 | Mouse | qPCR | ACCGAAGCTGAGAGCAACAT |
| *Tubb5* R | Mouse | qPCR | CACCATTTACCCCCAATGAG |
| *Atoh1* F | Mouse | qPCR | AATGACCACCATCACCTTCG |
| *Atoh1* R | Mouse | qPCR | TGTGGGATCTGGGAGATGTT |
| *Nfib* F | Mouse | qPCR | GTGTTCAGCCACACCACATC |
| *Nfib* R | Mouse | qPCR | GAGGATTCTTGGCAGGATCA |
| *Gabra1* F | Mouse | qPCR | CCAAGTCTCCTTCTGGCTCA |
| *Gabra1* R | Mouse | qPCR | CGGTTCTATGGTCGCACTTT |
| *Gabra6* F | Mouse | qPCR | TCTCCCCTGGCTCTTCATTA |
| *Gabra6* R | Mouse | qPCR | AAGTCTGGCGGAAGAAAACA |
| *Gli2* F | Mouse | qPCR | TCTACATGCCTTGGGTTGCTGTGGA |
| *Gli2* R | Mouse | qPCR | GAAAGTGTGGAGGCTGCAGGTGGTC |
| *Myc* F | Mouse | qPCR | TCGCTGCTGTCCTCCGAGTCC |
| *Myc* R | Mouse | qPCR | GGTTTGCCTCTTCTCCACAGAC |
| *Vglut1* F | Mouse | qPCR | CTGAGGAGGAGCGCAAATAC |
| *Vglut1* R | Mouse | qPCR | CAACGATGATGGCATAGACG |
| *Olfm3* F | Mouse | qPCR | CAACGGGTCCTCAGCTTAGA |
| *Olfm3 R* | Mouse | qPCR | CCGTCATCCAAGCACCAAAT |
